# Investigation of Zur-regulated metal transport systems reveals an unexpected role of pyochelin in zinc homeostasis

**DOI:** 10.1101/2024.01.07.574578

**Authors:** Valerio Secli, Emma Michetti, Francesca Pacello, Federico Iacovelli, Mattia Falconi, Maria Luisa Astolfi, Daniela Visaggio, Paolo Visca, Serena Ammendola, Andrea Battistoni

**Author notes:** Corresponding Author: (AB).

## Abstract

Limiting the availability of transition metals at infection sites serves as a critical defense mechanism employed by the innate immune system to combat microbial infections. *Pseudomonas aeruginosa* exhibits a remarkable ability to thrive in zinc-deficient environments, which is facilitated by intricate cellular responses governed by numerous genes regulated by the zinc-responsive transcription factor Zur. Many of these genes have unknown functions, including those within the predicted *PA2911-PA2914* and *PA4063-PA4066* operons. A bioinformatic analysis revealed that *PA2911-PA2914* comprises a TonB-dependent outer membrane receptor and an inner membrane ABC-permease responsible for importing metal-chelating molecules, whereas *PA4063-PA4066* contains genes encoding a MacB transporter, likely involved in the export of large molecules. Molecular genetics and biochemical experiments, feeding assays, and intracellular metal content measurements demonstrated that *PA2911-PA2914* and *PA4063-PA4066* are engaged in the import and export of the pyochelin-cobalt complex, respectively. Notably, cobalt can reduce zinc demand and promote the growth of *P. aeruginosa* strains unable to import zinc, highlighting pyochelin-mediated cobalt import as a novel bacterial strategy to counteract zinc deficiency. These results unveil an unexpected role for pyochelin in zinc homeostasis and challenge the traditional view of this metallophore exclusively as an iron transporter.

## Introduction

Zinc (Zn) is an essential micronutrient for all living organisms, serving as a structural or catalytic cofactor in numerous proteins involved in fundamental biological processes. In prokaryotes, Zn-binding proteins constitute approximately 6% of the proteome, including proteins involved in central metabolism, DNA replication and repair, antibiotic resistance, and virulence pathways (1,2). The intracellular concentration of this ion is finely regulated in cells to prevent problems due to imbalances. Indeed, Zn deficiency can lead to the loss of functionality of Zn-dependent proteins, whereas excess Zn can have toxic effects, including aberrant interaction with proteins and cellular components and competition with other relevant metal ions (3,4).

The importance of Zn is underscored by the concept of “nutritional immunity”, where vertebrates modulate the bioavailability of Zn and other transition metals at the host-pathogen interface to combat bacterial infections (5). As part of this defense mechanism, macrophages can accumulate Zn in phagosomes to poison the internalized pathogens with high concentrations of the metal (6,7). Conversely, serum or mucosal surfaces deplete Zn to hinder microbial growth during infections. This depletion occurs through metal redistribution among tissues and the action of metal-chelating proteins, such as the calprotectin released by neutrophils (8,9)

Pathogens have evolved adaptive mechanisms to acquire and maintain sufficient Zn levels to circumvent nutritional immunity. For instance, the high-affinity Zn-import system ZnuABC, a transporter of the ABC family widely conserved across many bacterial species, plays a crucial role in counteracting Zn deficiency by actively translocating Zn into the bacterial cytoplasm (2,10). Studies carried out in many bacteria, including *Salmonella enterica*, uropathogenic *Escherichia coli* (UPEC), and *Brucella abortus* have demonstrated a remarkable loss of pathogenicity when this system is inactivated (11–13).

Certain pathogenic bacteria employ additional Zn acquisition strategies, including synthesizing and releasing low molecular weight metal-binding molecules (metallophores) that act as extracellular metal chelators, facilitating selective Zn import. An example is represented by *Yersinia pestis*, where the inactivation of ZnuABC does not cause an evident loss of virulenceas the absence of this transporter is compensated by the ability of the siderophore yersiniabactin to also transport Zn (14). Zn-specific metal-binding metallophores, often called zincophores, have been identified in *Staphylococcus aureus* (15)*, Pseudomonas aeruginosa* (16), and a few other bacterial species (17) and have been proven to contribute significantly to the ability of these bacteria to cause infection (18,19). As an example, in *P. aeruginosa* the genes located in the *zrmABCD* operon, also known as cntOLMI (16) are involved in the synthesis and selective transport of pseudopaline, a metallophore mediating Zn import. This operon includes the genes encoding for the TonB-dependent transporter ZrmA, the nicotianamine synthase ZrmB, the opine dehydrogenase ZrmC and the inner membrane exporter ZrmD (16).

*P. aeruginosa* is a Gram-negative opportunistic pathogen responsible for infections that can be particularly threatening for immunocompromised individuals (20). In fact, its metabolic versatility, intrinsic resistance to many antibiotics, and ability to form biofilm, favor its rapid adaptation in the host environment, where it can give rise to both acute and chronic infections that are very difficult to treat*. P. aeruginosa* is particularly adept at responding to Zn fluctuation. It can withstand Zn deficiency conditions through the high-affinity Zn import system ZnuABC and the zincophore pseudopaline, which is synthesized and released in the Zn-limited environments characteristic of host tissues during infections (19). The functionality of these Zn import systems is crucial for *P. aeruginosa* to express a wide range of virulence features under conditions of Zn deficiency, allowing it to cause lung and systemic infections (19,21,22). Reduced Zn uptake affects the ability of *P. aeruginosa* to release extracellular Zn-dependent proteases, impairs swarming and swimming motility, hinders the synthesis of the exopolysaccharide alginate and biofilm formation, and reduces the release of the siderophore pyoverdine (22). Additionally, Zn-binding metallophores can interfere with the ability of calprotectin to sequester Zn, enhancing *P.aeruginosa* resistance to the nutritional immunity of the host (23). Moreover, *P. aeruginosa* produces and releases several metallophores, most of which have been characterized for their ability to deliver iron (Fe) to the cell (24). Even if these molecules are primarily involved in Fe transport, it should be remembered that they can strongly chelate other metals, as previously documented for the siderophore pyochelin (PCH) (25,26).

*P. aeruginosa* response to Zn deficiency is transcriptionally regulated by the Zn-uptake regulator protein Zur, which senses intracellular Zn levels and modulates the expression of its target genes (27,28). Among the genes regulated by Zur, there are those included in the *znuABC* and *zmrABCD* operons, responsible for the production of the previously mentioned Zn import systems Numerous genome-wide gene expression studies have identified Zur-regulated genes between the most highly expressed genes in *P. aeruginosa* recovered from infected tissues or cultured under host-mimicking conditions. These genes are overexpressed in various infection scenarios, underscoring their importance for the ability of *P. aeruginosa* to colonize the host environment (19,21–26).

Notably, the *P. aeruginosa* Zur regulon encompasses a larger number of genes than other bacteria and includes genes unique to this organism and whose role is not yet known (19,29). Given the high expression of many of these orphan genes during infection, understanding their role may foster our understanding of the interaction between *P. aeruginosa* and its host. This study explores the role of two uncharacterized Zur-regulated gene clusters, *PA2911-2914* and *PA4063-4066*, annotated as operons encoding putative metal import systems. The reported results demonstrate that these operons encode transport systems responsible for the import and export of the PCH-cobalt complex. Such results indicate that *P. aeruginosa* can adapt to Zn deficiency by substituting this metal with cobalt and reveal an unexpected role for PCH in Zn homeostasis, highlighting its multifunctional role beyond Fe transport.

## Results

### The *PA2911-2914* and *PA4063-4066* operons are regulated by Zn availability

The experiments reported in this paper were performed with the *P. aeruginosa* strain PA14. However, for ease of comparison with the extensive body of literature reporting data on these predicted operons, we will adopt the nomenclature of the reference strain PAO1. Consequently, we will refer to the operon *PA14_26420-PA14_26360* as *PA2911-2914* and to the operon *PA14_11320-PA14_11280* as *PA4063-PA4066* (Fig 1).

**Fig 1.**
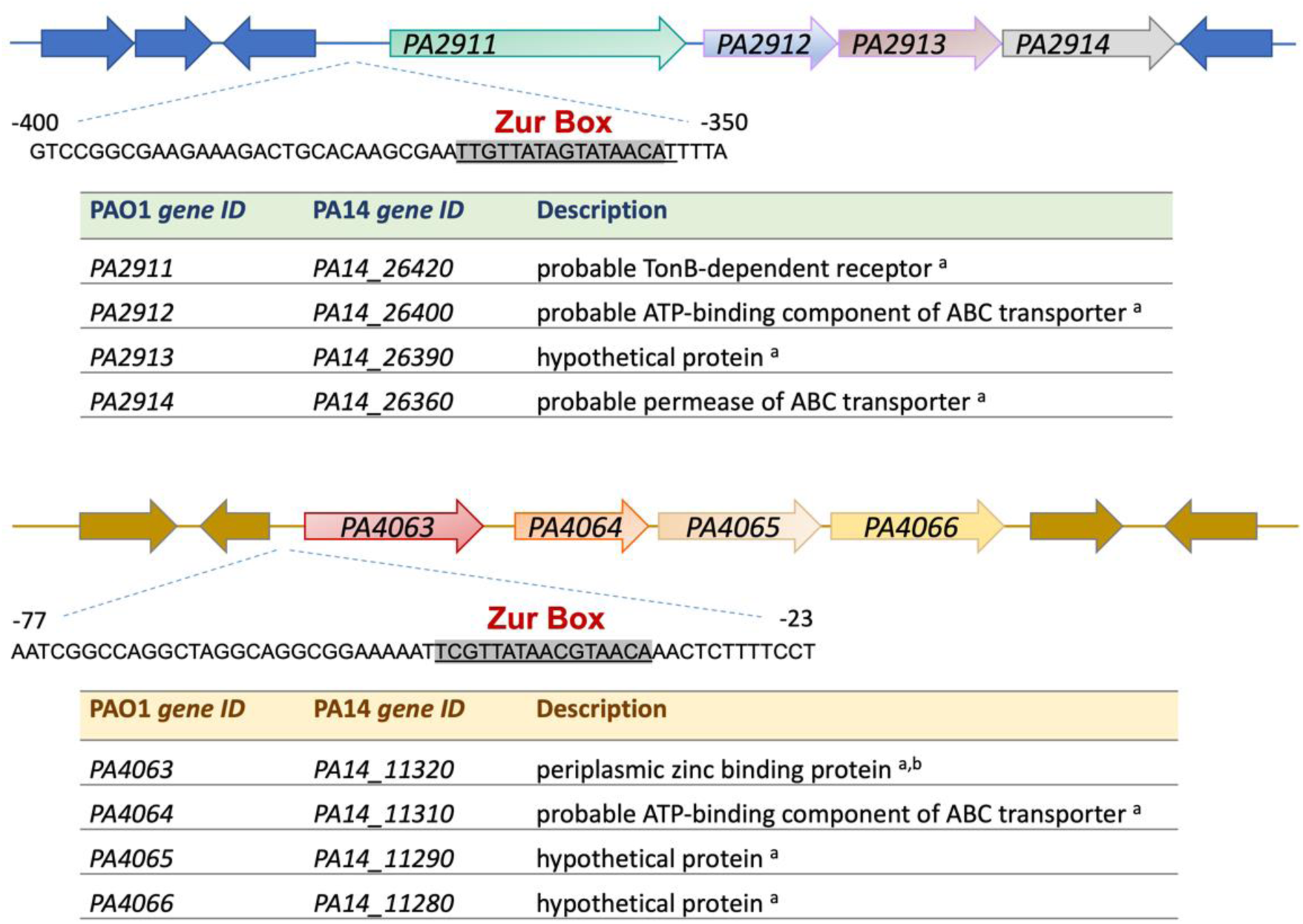
Schematic representation of the *PA2911-PA2914* and PA4063-PA4066 *operons*. The corresponding gene ID in PA14 strain and a brief description of the function are reported for each ORF. Sources are: ^a^ Pseudomonas Genome Database (https://www.pseudomonas.com) and ^b^(30). The Zur boxes (highlighted sequences) were previously identified by Pederick *et al.* (31).

The Zn-dependent regulation of *PA2911-2914* and *PA4063-4066* was initially assessed by comparing the relative abundance of the *PA2911* and *PA4063* transcripts (the first ORFs of each operon) in wild-type PA14 and in the *znuAzrmB* double mutant strain, both grown in E-VBMM with or without the addition of 3 µM ZnSO_4_. As shown in Figure 2A, *PA2911* and *PA4063* are significantly repressed in Zn-replete conditions (Fig 2A, wt + Zn and *znuAzrmB* + Zn bars), in agreement with the anticipated Zur-dependent regulation of these operons. Under Zn-deficient conditions, the expression of *PA4063* is comparable in wild-type PA14 and in the *znuAzrmB* double mutant strain. Conversely, the expression of *PA2911* significantly increases in the *znuAzrmB* background, where the intracellular concentration of Zn is lower than in the wild type (Fig. 2A, *znuAzrmB* bar). This observation implies a graded response of different Zur-regulated genes depending on the intracellular Zn availability. To investigate the role of *PA2911-PA2914* and *PA4063-PA4066* in Zn homeostasis, we deleted either *PA2914* or *PA4065*, the predicted inner membrane permeases of each operon, in both the PA14 wild-type and the *znuAzrmB* double mutant backgrounds. The growth curves of the resulting mutants (*PA2914, PA4065, PA2914znuAzrmB,* and *PA4065znuAzrmB*) were compared to their respective parent strains (i.e., wild-type and *znuAzrmB*) under Zn-restricted conditions. Figure 2B shows that neither *PA2914* nor *PA4065* inactivation impairs bacterial growth. However, when either *PA2914* or *PA4065* mutations are introduced into the *znuAzrmB* mutant strain, which lacks crucial Zn uptake systems, a marked decrease in bacterial growth is observed under low Zn availability conditions compared to a *znuAzrmB* mutant strain. The simultaneous deletion of *PA4065* and *PA2914* in the *znuAzrmB* background (resulting in the mutant strain *PA2914PA4065znuAzrmB*) does not cause any growth difference compared to each triple mutant (S1 Fig.).

**Fig 2.**
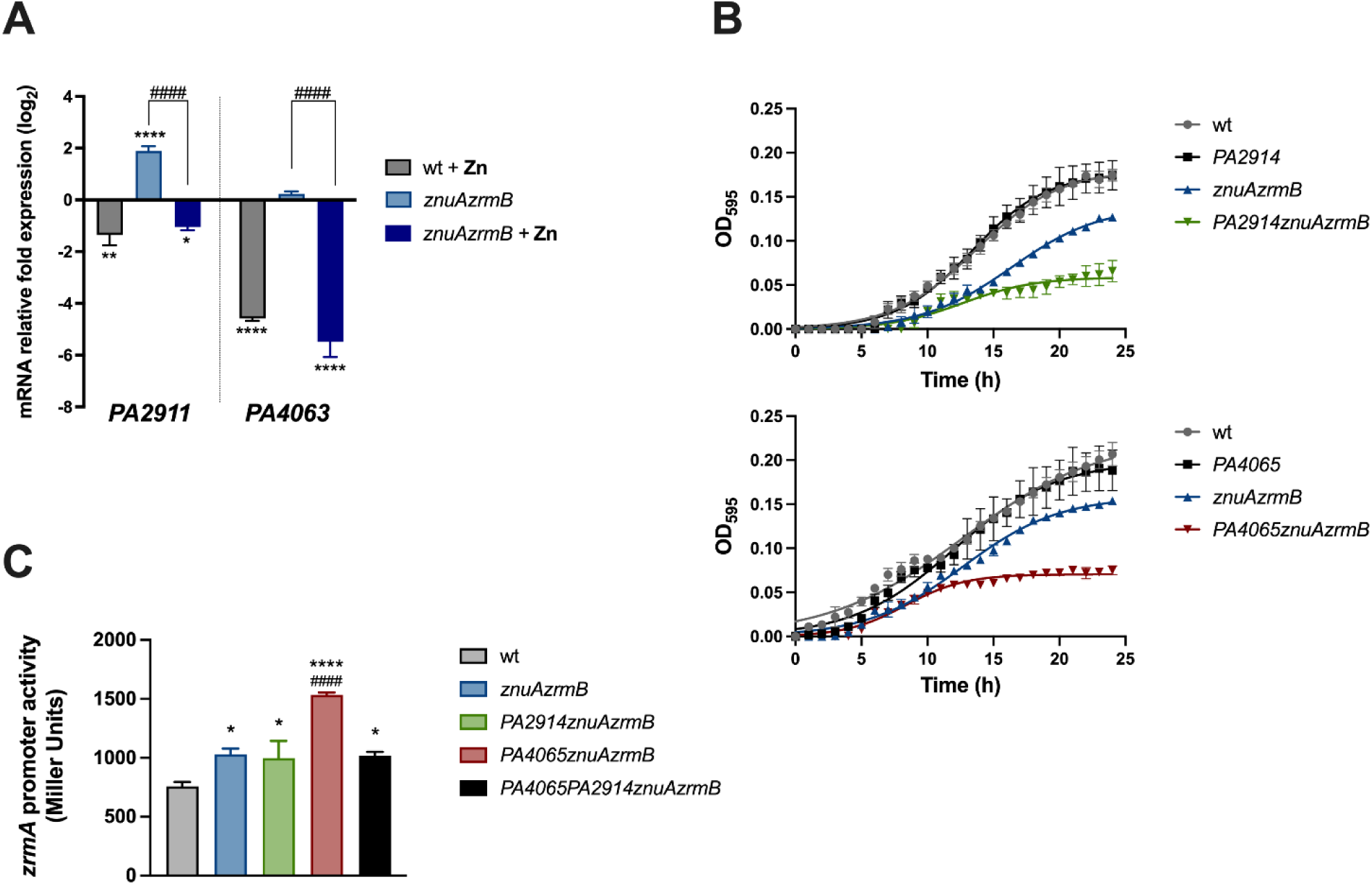
(A) Transcription of *PA4063* and *PA2911* in response to Zn availability. qRT-PCR on *PA2911* and *PA4063* from wild-type and *znuAzrmB* mutant strains grown in E-VBMM with or without ZnSO_4_ 3 μM, as indicated in the legend. Data are mean values ± S.D. of triplicates and expressed as Relative fold expression (log_2_ΔΔCt) compared to the gene expression in the wild-type strain grown in E-VBMM (control). Statistical analyses were performed by Two-way ANOVA and Bonferroni’s multiple comparison test. Asterisks indicate statistically significant differences between all samples and the control (*p<0.05; **p<0.005; ****p < 0.0001); hash signs indicate statistically significant differences between the *znuAzrmB* strain grown in E-VBMM and E-VBMM + Zn (^####^p < 0.0001). **(B) Contribution of *PA2914* and *PA4065* to PA14 growth under conditions of Zn limitation.** Growth curves of wild-type, *znuAzrmB,* and mutant strains carrying the *PA2914* deletion or the *PA4065* deletion, as indicated in the legends, grown in E-VBMM. Each symbol indicates the mean ± SD of triplicates, and lines represent nonlinear fit according to the Logistic Growth equation. **(C) Effect of *PA4065* and *PA2914* deletions on intracellular Zn request.** The beta-galactosidase activity driven by the *zrmA* promoter was evaluated after 20 hours of growth in E-VBMM in strains carrying plasmid *pzrmA*pTZ110. Each bar represents the mean ± SD of three biological replicates. Statistical significance was calculated by one-way ANOVA and Bonferroni’s multiple comparison test. Asterisks indicate statistical significance between wild-type (wt) and mutant strains (* p < 0.05; **** p < 0.0001); hash signs indicate statistical significance between *PA4065znuAzrmB* and the other mutant strains (^####^ p < 0.0001).

Next, since *zrmA* expression is significantly influenced by intracellular variations in Zn concentration (19), we employed the Zn-responsive *zrmA* promoter to assess whether the inactivation of *PA2914* or *PA4065* could alter the intracellular Zn status in bacteria grown in Zn-limited conditions. For this purpose, plasmid *pzrmA*pTZ110 (19) was introduced into wild-type PA14 and a panel of mutant strains with different combinations of mutations. The activity of the *zrmA* promoter was assessed in cultures grown in Zn-restricted conditions. The results presented in Figure 2C confirm that *zrmA* promoter activity is higher in the *znuAzrmB* strain than in the wild-type strain (19). Promoter activity increases even further in the *PA4065znuAzrmB* mutant, suggesting that the absence of *PA4065* intensifies the intracellular demand for Zn. In contrast, there are no differences in *zrmA* expression in the *PA2914znuAzrmB* and *znuAzrmB* strains. Intriguingly, the simultaneous deletion of *PA2914* and *PA4065* in a *znuAzrmB* background restores the *zrmA* promoter activity to the same level found in the *znuAzrmB* strain.

In summary, the results just described suggest that both *PA2911-PA2914* and *PA4063-PA4066* gene clusters are negatively regulated by Zn and contribute to the ability of *P. aeruginosa* to adapt to a Zn-limited environment. While the observed impact of simultaneous inactivation of both operons on the modulation of the Zur-responsive gene *zrmA* suggests a functional relationship between the two predicted operons, *PA4065* knock-out induces a stronger response of the *zrmA* promoter compared to the *PA2914* knock-out.

### *PA4064* and *PA4065* encode for a MacB-type exporter

A structural bioinformatics analysis was conducted to predict the functions of the proteins encoded by the *PA2911-2914* and *PA4063-PA4066* operons. Using the SWISS-MODEL web server, it was predicted that PA2911 serves as a TonB-dependent outer membrane receptor. At the same time, the *PA2912-PA2914* genes collectively encode an ABC family import system. Notably, all four components of this operon exhibit significant similarity to proteins involved in siderophore-mediated Fe uptake.

In addition, AlphaFold 2 was employed to predict the three-dimensional structures of PA4064 and PA4065. The analysis suggested that the two proteins form a complex with a fold characteristic of a MacB transporter, which is a class of ABC transporter that collaborates with other adaptor proteins to facilitate the efflux of antibiotics, toxins, or siderophores out of the bacterial cell (32). MacB transporters generally consist of two interacting MacB monomers, each containing four transmembrane helices, an N-terminal nucleotide-binding (NBD) domain, and a long periplasmic domain between TM1 and TM2. In this case, the putative MacB transporter predicted by Alphafold 2 (Fig 3) is formed by the assembly of two pairs of subunits. *PA4064* indeed codes for the N-terminal nucleotide-binding domains (Fig 3, green and orange), while *PA4065* codes for the TM helices and the periplasmic domains (Fig 3, gray and red). The pLDDT values for both domains are notably high (S2 Fig.), indicating the high quality and reliability of the models. These findings suggest that the *PA4063-4066* operon does not encode for a metal import system but, more plausibly, for a transporter involved in the export of a molecule contributing to cellular responses to Zn deficiency.

**Fig 3.**
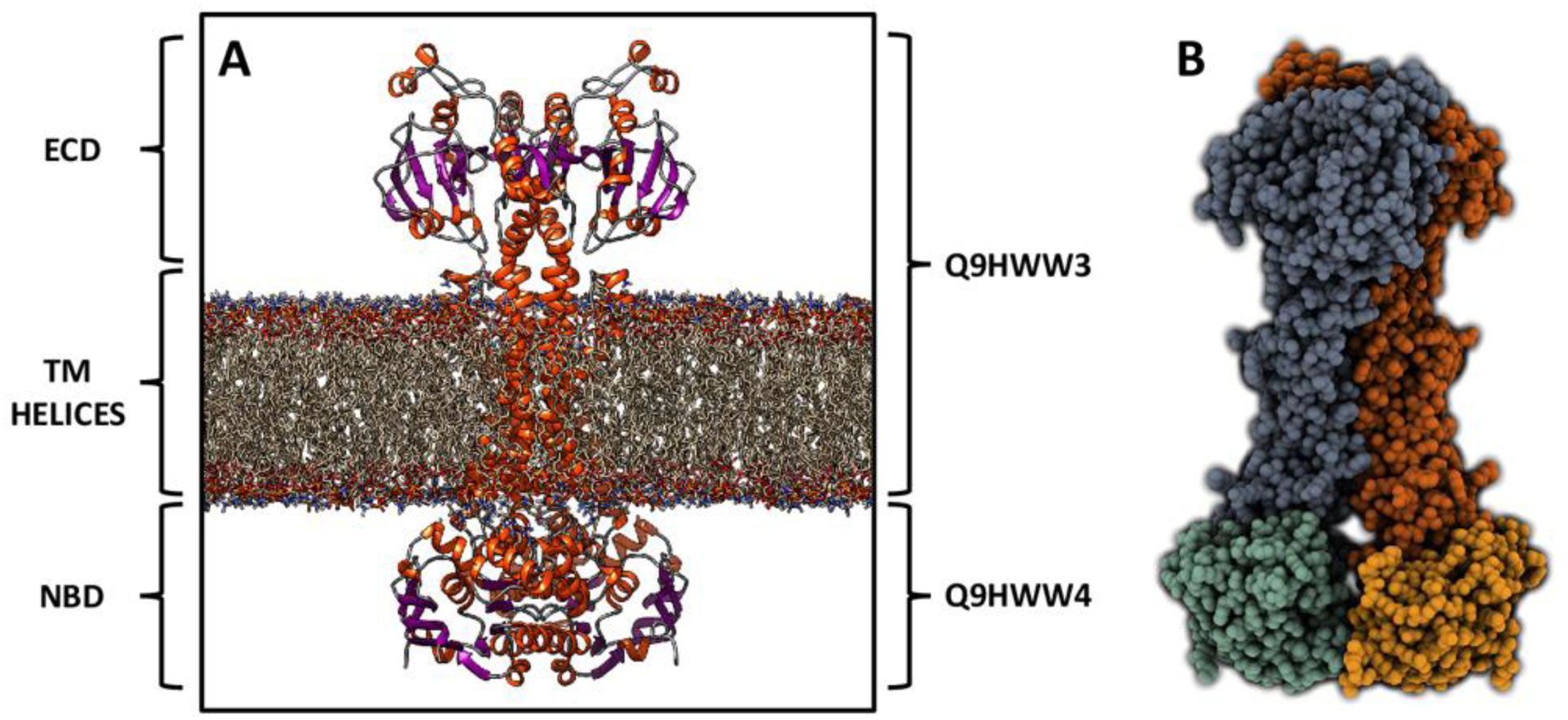
Molecular modelling of PA4064-PA4065. Molecular representation of the predicted MacB transporter embedded in a membrane. α-helices are represented as orange spirals, β-strands through violet arrows, while the membrane is shown using a stick representation.

### Pyochelin production is downregulated in the *znuAzrmB* mutant strain

Differences in the pigmentation of liquid cultures between the wild-type and the *znuAzrmB* strain of *P. aeruginosa* PA14 grown overnight under Zn-limiting conditions (VBMM) were previously reported (22). Notably, the wild-type strain showed a greenish tint, while the *znuAzrmB* cultures were characterized by a pale blue color. This difference was correlated with a decrease in pyoverdine (PVD) production (22). In this study, we observed that the inactivation of either *PA2914* or *PA4065* in the *znuAzrmB* strain caused the cultures to regain a color very similar to that of the wild-type PA14 (S3 Fig).

Besides PVD, which is produced in low amounts by the PA14 strain in these conditions, *P. aeruginosa* secretes other pigments that absorb light in the UV-visible spectrum. Therefore, we decided to investigate whether these phenotypic differences could be attributed to changes in the abundance of one of these metabolites by analyzing the aqueous and ethyl acetate fractions obtained from cell-free supernatant. Fluorimetric studies on the aqueous fraction confirmed the reduction of PVD secretion in the *znuAzrm*B mutant previously observed in the *znuAzrmA* mutant strain (22). Even if differences in PVD production can contribute to the different colors of bacterial cultures, the ethyl acetate fractions, which contain pigments less hydrophilic than PVD, exhibited much more marked differences.

Albeit with small quantitative differences, the UV-visible spectra of the ethyl acetate fractions from the wild-type, *PA2911znuAzrmB,* and *PA4065znuAzrmB* strains all displayed three major peaks around 250, 315, and 370 nm (Fig 4A). In contrast, the *znuAzrmB* sample showed only a weak peak around 315 nm. Since the absorption spectrum of purified apo-PCH is characterized by two major peaks at 248 and 313 nm (33), we hypothesized that the color changes among the different supernatants could be largely due to differences in PCH secretion. The peak at 370 nm is likely due to residual pyocyanin contamination (34). Electro Spray Ionization Mass Spectrometry (ESI-MS) analyses of the ethyl acetate fractions from the wild-type, *PA4065znuAzrmB* and *PA2914znuAzrmB* strains confirmed the presence of PCH, but failed to detect PCH in the *znuAzrmB* sample (S4 Fig.). S4 Fig. also reveals that the mass spectrum profile of the supernatant from the *znuAzrmB* double mutant is extremely simplified compared to the other strains and that in addition to PCH, other metabolites are absent or strongly reduced.

**Fig 4.**
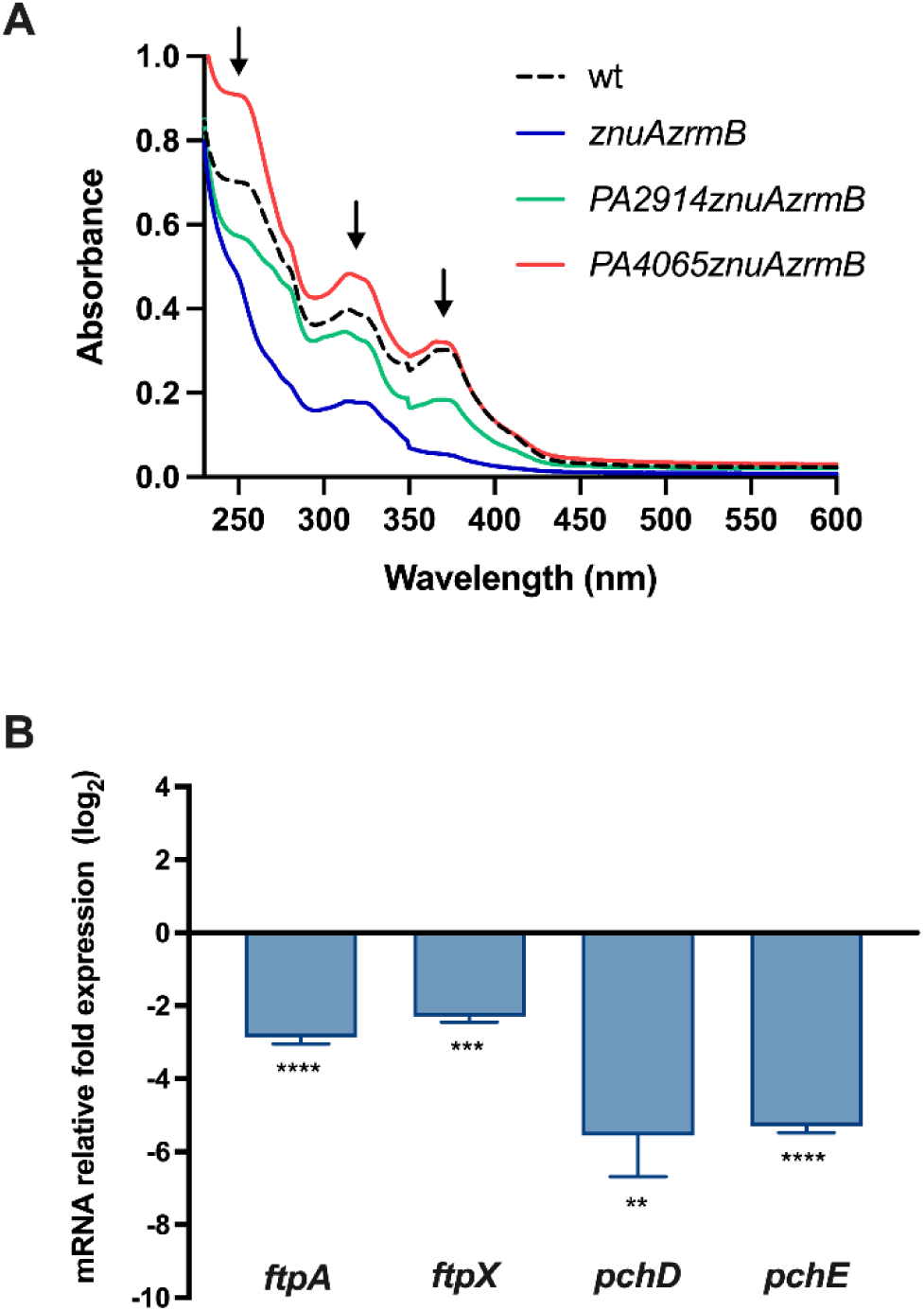
PCH synthesis is downregulated in the *znuAzrmB* strain. **(A)** UV-vis absorbance spectra (230 - 600 nm) of ethyl acetate fractions extracted from the culture supernatants. The arrows denote peaks at 250 and 315 nm wavelength matching the PCH spectrum. **(B)** Relative fold expression (log_2_ΔΔCt) of genes involved in PCH uptake (*ftpA* and *fptX*), and synthesis (*pchD* and *pchE*), in the *znuAzrmB* mutant grown in E-VBMM, relative to the PA14 wild type. Data are mean values ± S.D. of triplicates, and statistical analysis was performed by Paired T test. Asterisks indicate statistically significant differences between the *znuAzrmB* strain and wild-type, grown in the same conditions (**p<0.005; ***p<0.0005; ****p < 0.0001).

To explore whether the difference in PCH secretion between wild-type and *znuAzrmB* strains might be related to a modulation of gene expression in PCH metabolic pathway, we conducted qRT-PCR analyses of the transcription levels of genes involved in PCH uptake (*fptA* and *fptX*) and synthesis (*pchD* and *pchE*) in wild-type and *znuAzrmB* strains grown in E-VBMM. Results indicated that the transcription of *pchD*, *pchE, fptA,* and *fptX* was significantly downregulated in the *znuAzrmB* strain compared to the wild-type (Fig 4B).

### The inability to export pyochelin through the MacB transporter increases the cellular demand for Zn

The results discussed in the previous section demonstrate a significant decrease in PCH production in the *znuAzrmB* mutant that is largely restored when the mutations in *znuA* and *zrmB* are combined with the deletion of *PA2914* or *PA4065* (S3 Fig.). Furthermore, based on activation of the of *zrmA* promoter, it was observed that the intracellular requirement for Zn is greater in the *PA4065znuAzrmB* triple mutant compared to the *znuAzrmB* double mutant. However, the further deletion of *PA2914* restores expression of *zrmA* to levels observed in the double mutant (see Figure 2C). These findings led us to hypothesize that *PA2911-PA2914* and *PA4063-PA4066* operons are involved in the transport of PCH in and out of the cell. This mechanism could be related to bacterial adaptation when facing severe Zn limitation.

To explore this hypothesis, we analyzed the transcriptional activity of *zrmA* in the mutant strains *znuAzrmB*, *PA2914znuAzrmB,* and *PA4065znuAzrmB* compared with each isogenic *pchE* mutant (*znuAzrmBpchE*, *PA2914znuAzrmBpchE,* and *PA4065znuAzrmBpchE)*, grown in E-VBMM. The *pchE* strains cannot produce mature PCH due to the inactivation of one of the enzymes involved in the biosynthetic process. Figure 5A shows that the *zrmA* promoter is expressed at higher levels in the *znuAzrmBpchE* strain than in the *znuAzrmB* strain, suggesting that impairment of PCH synthesis enhances the intracellular Zn demand. No changes in transcription were observed when the *pchE* and *PA2914* mutations were combined, consistent with the hypothesis that the *PA2911*-*2914* operon encodes a PCH import system. Remarkably, *zrmA* transcription is restored to levels seen in the *znuAzrmB* strain when the deletion in *pchE* is introduced in the *PA4065znuAzrmB* mutant strain, indicating a role for PCH in the high activity of the *zrmA* promoter in the triple mutant. No changes in transcription were observed when the *pchE* and *PA2914* mutations were combined, suggesting that the *PA2911-2914* encodes a PCH import system.

**Fig 5.**
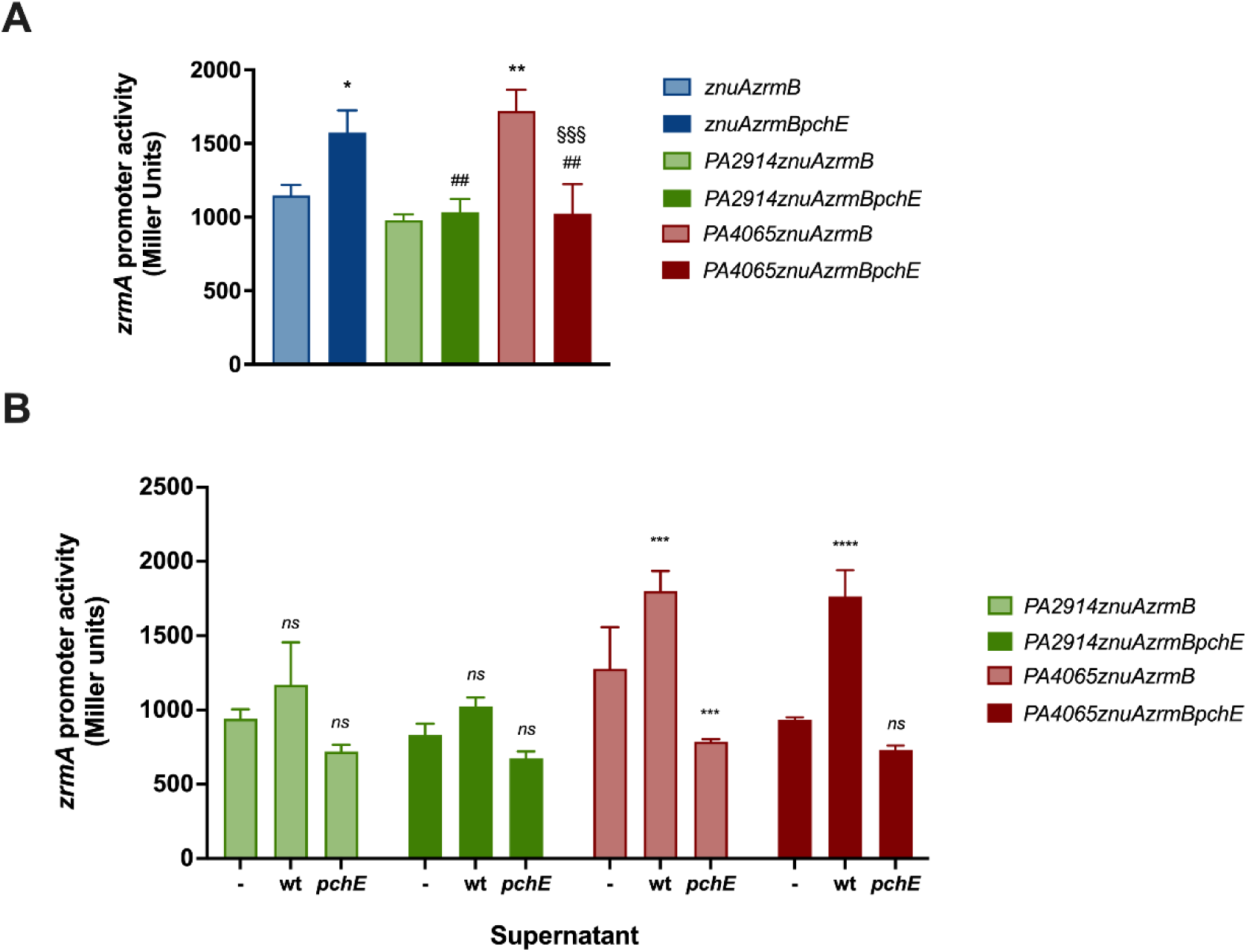
Effects of PCH on intracellular Zn request. **(A)** The *zrmA* promoter activity was evaluated in strains carrying plasmid *pzrmA*pTZ110, grown in E-VBMM for 20 hours. Each bar represents the mean ± SD of three biological replicates, and statistical significance was calculated by one-way ANOVA and Bonferroni’s multiple comparison test. Asterisks indicate the statistical difference with the *znuAzrmB* strain (* p < 0.05; ** p < 0.01); hash signs indicate the statistical difference between the *znuAzrmBpchE* and the other strains (^##^ p < 0.01); § show statistical difference with the *PA4065znuAzrmB* strain (^§§§^ p < 0.005) **(B)** The *zrmA* promoter activity was evaluated in strains carrying plasmid *pzrmA*pTZ110, grown for 20 hours in E-VBMM (-) or supplemented with the ethyl acetate fraction of supernatants from wild-type (wt) or *pchE* strains (*pchE*). The decrease in *zrmA* expressions in cultures treated with supernatants from the *pchE* strain is likely caused by trace amounts of Zn introduced during the ethyl acetate extraction. Data represent means ± SD of three biological replicates. Two-way ANOVA and Tukey’s multiple comparison tests were performed to assess significant differences (*** p < 0.001; **** p < 0.0001; *ns*: not significant)

The impact of PCH on the Zn intracellular status was then analyzed by assessing the activity of the *zrmA* promoter in *PA2914znuAzrmB*, *PA2914znuAzrmBpchE*, *PA4065znuAzrmB,* and *PA4065znuAzrmBpchE* strains, grown in VBMM supplemented with the ethyl acetate fraction from wild-type or *pchE* strain. As shown in Figure 5B, no significant differences were observed among the *PA2914* defective strains, indicating that the disruption of this importer has no impact on intracellular Zn availability, either in the presence or absence of PCH in the growth medium. In contrast, feeding with a PCH-containing supernatant induces the *zrmA* promoter in strains lacking *PA4065* (Figure 5B). These results suggest that the increased expression of *zrmA* in the *PA4065znuAzrmB* mutant may be attributed to the intracellular accumulation of PCH, causing dysregulation of Zn homeostasis, and that PA4065 plays a role in the export of PCH.

### PA2914 and PA4065 are involved in pyochelin trafficking

To test the hypothesis that PA4065 is involved in PCH export, we used a luminescent biosensor (35) to measure the concentration of PCH in cell-free supernatants and cell lysates from the wild-type, *znuAzrmB*, *PA4065* and *PA4065znuAzrmB* strains (Fig 6). This analysis not only confirmed that PCH production is below the detection limit in the *znuAzrmB* strain but also revealed that the deletion of *PA4065* significantly enhances the intracellular accumulation of PCH, particularly in the *PA4065znuAzrm*B triple mutant. PCH secretion was comparable among the other strains, with a slight yet significant increase observed in the *PA4065znuAzrmB* supernatant. The ability of mutants lacking PA4065 to export PCH and the increase of extracellular PCH in the triple mutant, which appears to be correlated with the high concentration of intracellular PCH in this strain, confirms the existence of PCH export mechanisms independent from the PA4064-PA4065 MacB exporter identified in this study.

**Fig 6.**
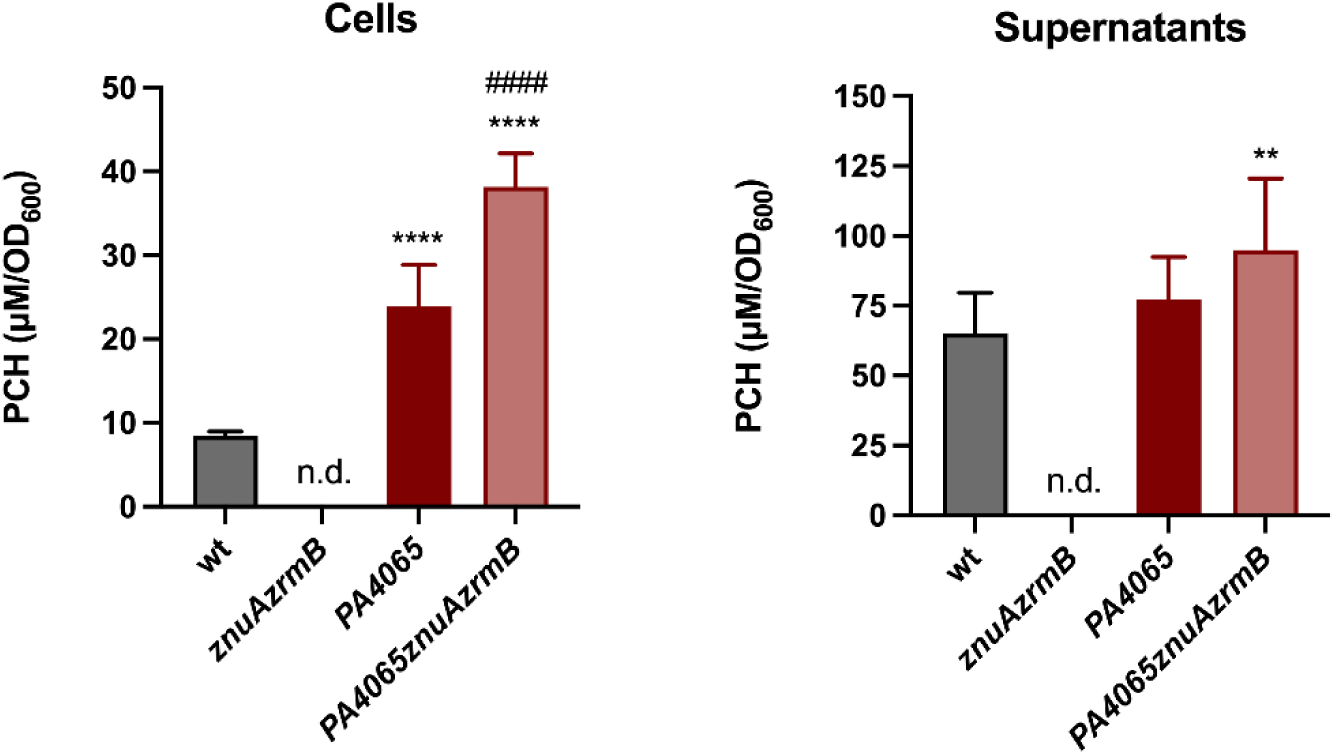
Quantification of PCH in cell lysates and cell-free supernatants. Wild-type and mutant strains were grown in VBMM, and PCH production was quantified using the *P. aeruginosa ΔpvdAΔpchDΔfpvA PpchE*::*lux* biosensor, as described in Materials and Methods. Data are expressed as μM/OD600; each bar is the mean value of three independent experiments ± SD. One-way ANOVA and Tukey’s multiple comparison tests were performed to assess significant differences among wild-type (wt) and mutant strains (** p < 0.01; **** p < 0.0001) and between *PA4065* and *PA4065znuAzrmB* (#### p < 0.0001). PCH was not detected in the *znuAzrmB* strain samples (n.d.)

The influence of PCH on intracellular Zn homeostasis was evaluated by supplementing the growth medium with purified apo-PCH and analyzing the *zrmA* promoter activity in different mutant strains (Fig 7). Exogenous PCH significantly increases the Zn demand in all strains except the one lacking PA2914. This differential response of the various strains to incubation with PCH seems to exclude that the increase in *zrmA* expression is due to a chelating effect of PCH, whose ability to bind Zn is well known (26,36). Instead, this observation aligns with the hypothesis that PA2914 plays a role in PCH uptake.

**Fig 7.**
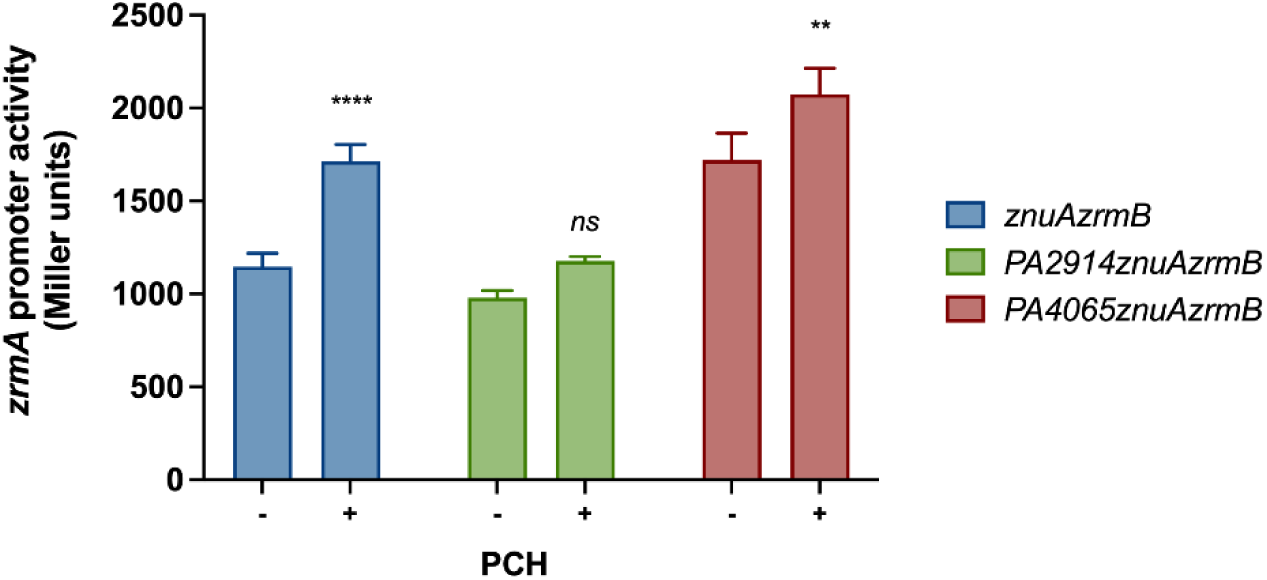
Influence of PCH on intracellular Zn request. The *zrmA* promoter activity was evaluated in *znuAzrmB* (left panel), *PA2914znuAzrmB* (middle panel), and *PA4065znuAzrmB* (right panel) strains carrying plasmid *pzrmA*pTZ110, grown in VBMM supplemented (+) or not (-) with 1 µM apo-PCH. Each bar is the mean value of three biological replicates ± SD. Two-way ANOVA and Sidak’s multiple comparison tests were performed to assess significant differences, and asterisks indicate significant differences between untreated and PCH-treated samples (**p < 0.01; ****p < 0.0001); *ns*: not significant.

### The inactivation of *PA4065* or *PA2914* has an impact on intracellular cobalt content

To investigate the hypothesis that *PA2911-PA2914* and *PA4063-PA4066* are involved in a PCH-mediated mechanism of metal acquisition aimed at counteracting Zn-starvation conditions, we measured the metal content in various mutant strains grown in E-VBMM supplemented with trace amounts of transition metals (19). Consistent with previous observations, the Zn content of the *znuAzrmB* mutant is significantly lower than that determined for both the wild-type and the *znuA* mutant strains, confirming the crucial role of pseudopaline in adapting to Zn-deficiency (19). The introduction of *PA2914*, *PA4065* or *pchE* deletions into the *znuAzrmB* strain did not further decrease cellular Zn content (Fig 8, left panel). This suggests that neither PCH nor PA4065 or PA2914 is directly involved in Zn uptake under conditions of metal restriction. In addition, no significant differences were observed in these strains with respect to metals such as copper (Cu), nickel (Ni), or manganese (Mn). While changes in Fe content were moderate, strains lacking *pchE* exhibited a trend towards decreased Fe content (S3 Table). In contrast, substantial differences were observed in cobalt (Co) content (Fig 8, right panel). While the inactivation of *znuA* does not affect Co content, strains unable to synthesize PCH or pseudopaline displayed a sharp decrease in Co accumulation. This result suggests that PCH and pseudopaline play a role in Co uptake during Zn starvation. Intriguingly, while the Co content of the *PA2914znuAzrmB* remains comparable to that of all the *pchE* mutants, the deletion of *PA4065* led to a significant and unexpected increase of Co content unless PCH production was abrogated. These findings strongly suggest that PA2914 and PA4065 are involved in the trafficking of the PCH-Co complex in and out of the cell during Zn starvation.

**Fig 8.**
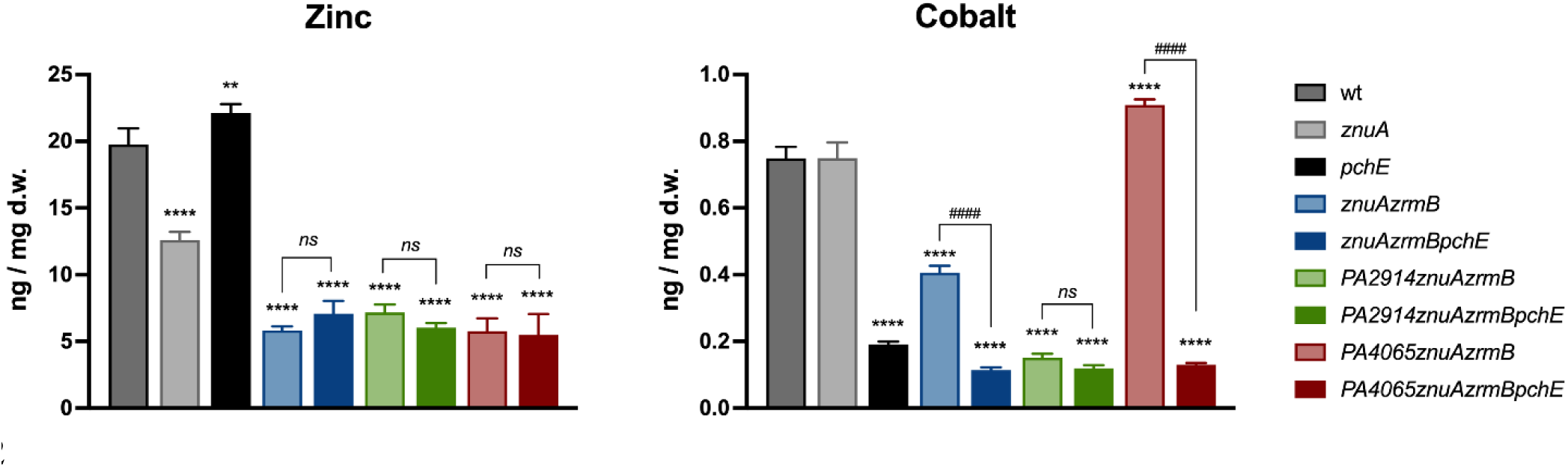
Intracellular Zn and Co content. ICP-MS analyses of bacteria grown in E-VBMM with trace metals (0.2 μM ZnSO_4_ and 0.1 μM FeSO_4_, NiSO_4_, Co(NO_3_)_2_, CuSO_4_ and MnCl_2_). Bars are the mean value of three biological replicates ± SD. Statistical analyses were performed by One-way ANOVA. Asterisks indicate statistically significant differences between mutant strains and wild-type (**p < 0.05; ****p < 0.0001); hash signs indicate pairwise statistically significant differences (^####^p < 0.0001; *ns*: not significant).

### The absence of PCH alleviates the cobalt-dependent growth impairment

The PA14 wild-type strain is sensitive to Co, which causes a growth impairment even at relatively low concentrations. In contrast, we have shown that Co promotes the growth of the *znuAzrmB* mutant, indicating that this metal can at least partially compensate for low Zn availability (19). To investigate the role of PCH in Co accumulation, we conducted growth experiments with the *PA4065znuAzrmB* and *PA4065znuAzrmBpchE* mutant strains in E-VBMM, with or without Co supplementation. As shown in Fig 9, Co supplementation caused a noticeable growth delay in the *PA4065znuAzrmB* mutant, similar to what was observed in the wild-type strain (S5 Fig.), likely explained by the PCH-mediated Co accumulation in the cell. In contrast, the quadruple mutant *PA4065znuAzrmBpchE*, where PCH biosynthesis is abolished, displays no growth defect when grown in the presence of Co. These findings provide additional evidence that PCH mediated Co accumulation and that the MacB exporter formed by PA4064-PA4065 serves as a PCH exporter. This mechanism appears to be valuable in preventing excessive intracellular Co accumulation.

**Fig 9.**
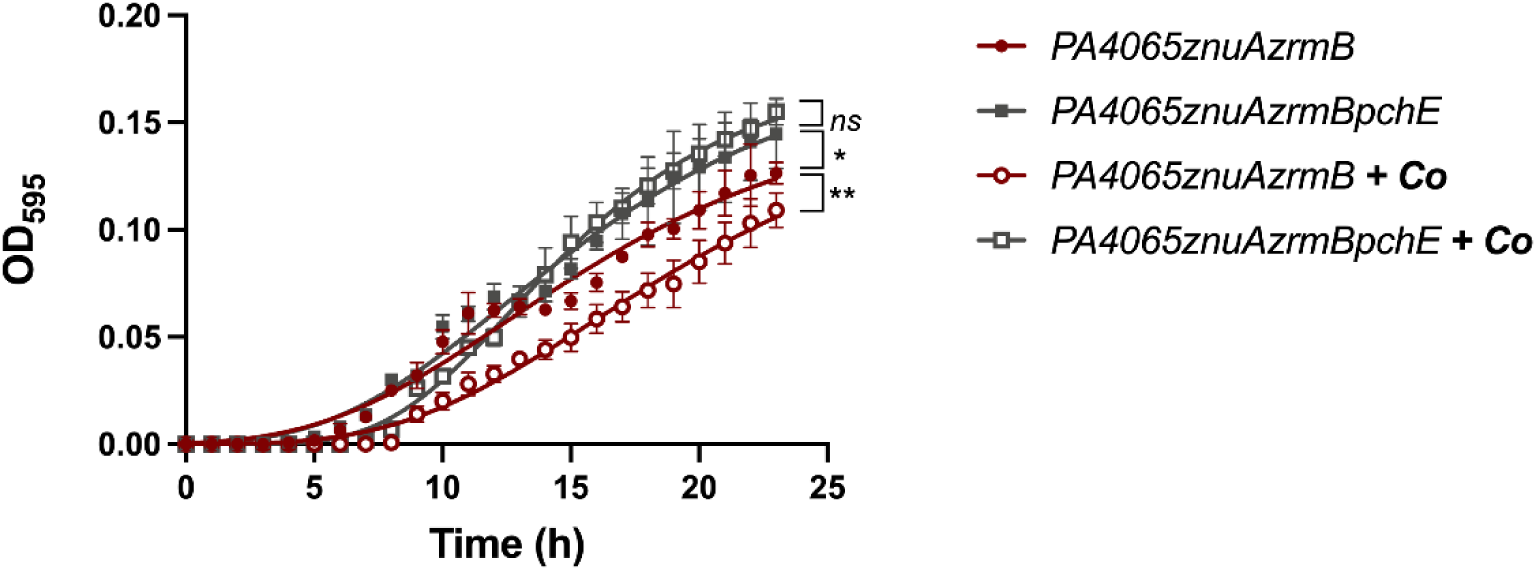
Effect of *pchE* on Co supplementation of strains carrying the *PA4065* deletion. Growth curves of PA4065*znuAzrmB* and *PA4065znuAzrmBpchE* strains in E-VBMM (filled symbols) and E-VBMM + Co(NO_3_)_2_ 10 μM (empty symbols). Each point indicates the mean value ± SD of triplicates, and lines represent nonlinear fit according to the Logistic Growth equation. Statistical differences among the end points of each curve were analyzed by two-way ANOVA (*p < 0.05; **p < 0.01; *ns*: not significant).

## Discussion

The ability to efficiently acquire Zn in environments where this metal is poorly available plays a crucial role in the interaction of *P. aeruginosa* with its hosts. This is demonstrated by the impaired expression of several virulence features and the marked loss of virulence of strains with reduced Zn acquisition ability (19,37). *P. aeruginosa* exhibits robust responses to Zn deficiency, as evidenced by its efficient proliferation in Zn-poor media and resistance to the antimicrobial activity of the Zn-sequestering protein calprotectin, even in the absence of a functional ZnuABC transporter (21,28). The identification of an extracellular metal capture mechanism based on the pseudopaline metallophore produced by the ZrmABCD system has provided evidence that *P. aeruginosa* uses additional systems to obtain Zn (16,19). However, our understanding of *P. aeruginosa* adaptations to Zn deficiency is still limited. Notably, the Zur operon in this species includes over 30 genes, many of which are yet to be characterized, and some of which are not found in closely related *Pseudomonas* species (29). For instance, the *zrmABCD* operon, involved in pseudopaline synthesis and transport, is highly conserved in all *P. aeruginosa* strains, including clinical isolates, but is not found in other closely related species colonizing soil, water, and plants such as *P. fluorescens*, *P. putida*, *P. stutzeri* and *P. syringae*. This suggests that pseudopaline could be particularly important for promoting Zn recruitment within animal hosts.

To shed light on the function of Zur-regulated genes of unknown function, this study aimed to investigate two previously uncharacterized Zur-regulated operons, *PA2911-2914* and *PA4063-4066*, both of which include genes encoding putative metal transporters (Fig. 1).

The results presented in Fig. 2 confirm that both operons are regulated by Zn. Mutants lacking *PA2911* or *PA4065* show similar growth to the wild-type strain in a Zn-poor medium. However, when these mutations were introduced into a strain lacking ZnuA and ZrmB, a significant reduction in growth was observed, highlighting the contribution of these two operons to adaptation under conditions of Zn deficiency, especially in bacteria strongly defective in Zn acquisition.

However, the results reported in Fig. 2C raise questions regarding the mechanisms by which these operons contribute to adaptation under conditions of Zn deficiency. Transcription arising from the *zrmA* promoter, that is sensitive to changes in intracellular Zn concentration, was assessed introducing the Zur-responsive *zrmA::lacZ* promoter fusion vector in different mutants. It was observed that *zrmA* expression is significantly higher in the *PA4065znuAzrmA* triple mutant than in the *znuAzrmA* double mutant. However, the expression of *zrmA* in the *PA2911PA4065znuAzrmA* quadruple mutant was indistinguishable from that observed in the *znuAzrmB* mutant, suggesting a functional relationship and compensation between the systems encoded by the two operons.

To find an interpretation of these results, we performed a bioinformatic analysis on the proteins encoded by the *PA2911-2914* and *PA4063-PA4065* operons. The *PA2911-2914* operon encodes a TonB-dependent outer membrane receptor (PA2911) and the components of a classical importer of the ABC family (PA2912-PA2914). These proteins show similarities with other transporters involved in the entry of siderophores or vitamin B12 into the cell, leading to the hypothesis that they constitute a system capable of mediating the transport of a metal bound to a metallophore from outside the cell to the cytoplasm. More unexpected were results deriving from the analysis of the *PA4063-4066* operon. Previous studies have annotated this system as an ABC transporter involved in Zn uptake (29,38). However, a protein prediction analysis carried out with Alphafold suggested that the PA4064 and PA4065 proteins are subunits of an exporter of the MacB family. MacB proteins represent an atypical family of ABC transporter described in many Gram-negative bacterial species, including *E. coli*. These transporters typically bind a MacA component and the inner membrane protein TolC to form a tripartite efflux pump. The tripartite complex MacA-MacB-TolC is a transmembrane machinery that extends both on the plasma membrane and on the outer membrane and actively extrudes large molecules, including macrolide antibiotics, virulence factors (e.g., pyoverdine in the case of *P. aeruginosa*), peptides and precursors of the cell envelope (39–42). The main function of the MacB transmembrane domain is to transmit conformational changes from one side of the membrane to the other, with a process defined mechanotransmission (39). While previously characterized MacB transporters were encoded by a single polypeptide chain, this *P. aeruginosa* MacB is formed by the association of two independent polypeptides, with PA4064 forming the ATP binding domain and PA4065 encoding the membrane channel.

Despite this difference in gene organization, the predicted structure of PA4064/PA4065 perfectly overlaps the structure of known MacB proteins. Although further studies are needed to clarify the export mechanism of this unique MacB transporter, it is important to note that this organization is conserved in other bacterial species. For example, a gene cluster analogous to *PA4063-PA4066* is present in *Vibrionaceae* and other bacteria, all under the control of Zur (43). The other two operon members, namely *PA4063* and *PA4066,* encode for soluble periplasmic proteins of known structure but still unclear function. PA4063 is a metal-binding protein able to bind Zn or Co ions at relatively low affinity, proposed to play a role as a periplasmic metal chaperone (30). In *Vibrio cholerae*, its homologous protein, ZrgA, is required for the proper function of the whole transport system (44). The crystal structure of PA4066 has revealed that this protein may assemble in a tetrameric structure with no evident ability to bind Zn. Although the function of PA4063 and PA4066 remains undisclosed, the observation that even in other bacteria the genes encoding their homologs are invariably associated with those for proteins like PA4064 and PA4065 corroborates their involvement in supporting the export functions of the MacB transporter.

The identification of PA4063-4066 as an export system led to the hypothesis that the results shown in Fig. 2C could be rationalized assuming that the PA2911-2914 uptake system is responsible for the internalization of a metallophore which must then be eliminated from the bacterial cytoplasm through the action of the MacB exporter coded by *PA4063-4066*. The intracellular accumulation of metal-binding molecules can be harmful to cells due to their ability to interfere with metal binding to proteins, which can cause a condition of apparent metal deficiency. For example, it has been shown that the intracellular accumulation of staphylopine (a molecule structurally related to pseudopaline) is toxic to *Staphylococcus aureus* (45,46). In line with this, it has been demonstrated that disruption of either *zrmD* or *PA4063* caused a dramatic loss of growth ability in Zn-devoid airway mucus secretion (47).

The next challenge was to identify the substrate imported by PA2911-2914. It could have been a still unknown metallophore, perhaps produced specifically in conditions of Zn deficiency, or an already known molecule with other associated functions. The results in Fig.4 showed a significant reduction in the release of PCH in the *znuAzrmB* mutant, stimulating further investigations of the potential involvement of this metal chelator in the response to Zn deficiency. PCH is well known to promote microbial growth by solubilizing ferric iron and accelerating metal transport within the cell (48). However, *in vitro,* PCH can bind other metals with high affinity, including Ag^+^, Co^2+^, Cu^2+^, Ni^2+^, Pb^2+,^ and Zn^2+^ (49). Moreover, Brandel *et al*. showed that that PCH strongly chelates divalent metals such as Zn (pZn = 11.8 at pH 7.4) forming 1: 2 (M2^+^/PCH) complexes (26). While the ability of PCH to bind Zn is remarkable, it does not mediate its import into cells. This is due to the transport selectivity of the FptA receptor, which efficiently mediates the uptake of the Fe-PCH complex, and, to a lesser extent, of complexes with Ni, Co, and gallium, but not those involving other metals (49,50). Interestingly, the ability of FptA to mediate entry of the PCH-Co complex affects the production of PCH (51). It is also worth mentioning some recent studies reporting that in *Yersinia pestis*, the yersiniabactin siderophore, structurally very similar to PCH, binds Zn^2+^ and transports it to the cell via the transporter YbtX (14,52). Likewise, the use of yesiniabactin as a zincophore has been documented in *Escherichia coli* (53).

These findings led to the hypothesis that *PA2911-2914* codes for an alternative PCH import system, while the MacB transporter encoded by *PA4064/PA4065* serves as an export pump for PCH. Spectrophotometric analyses of bacterial supernatants (Fig. 4), feeding assays and experiments with purified PCH (Fig. 5 and Fig. 7), and measurements of PCH concentration in bacterial extracts and supernatants (Fig. 6) support this hypothesis. Moreover, an ICP-MS analysis has revealed that no relevant differences in Zn concentration were observed between the *znuAzrmB* strain and the *PA2911znuAzrmB* and *PA4065znuAzrmB* mutants. However, the *PA4065znuAzrmB* mutant exhibited a disproportionate accumulation of Co, which was completely abolished in the *PA4065znuAzrmBpchE* mutant strain (Fig. 8). All these data converge to support the hypothesis that *PA2911-2914* and *PA4063-4066* gene clusters are involved, respectively, in the import and export of PCH. Specifically, the results indicate that PA2911-2914 is involved in the uptake of the PCH-Co complex while the MacB transporter is involved in the export of excess Co, possibly in complex with PCH. These observations, therefore, suggest that one of the strategies adopted by *P. aeruginosa* to respond to a condition of Zn deficiency is to activate a PCH-based system to supply the cell with Co. Importantly, ICP-MS data reveal that both PCH and pseudopaline participate in the acquisition of Co. Furthermore, the observation that *PA2911* is more expressed in the *znuAzrmB* mutant than in the wild-type strain supports the hypothesis that this transport system becomes necessary when other routes for Zn acquisition are already defective.

The import of Co under conditions of Zn deficiency does not appear to be a causality. Co has the same atomic radius as Zn and can be incorporated in many proteins instead of Zn, functionally replacing the original metal (54). Recent studies have revealed that in *Salmonella* Co supplementation can rescue the growth defects caused by Zn deficiency and that Co can overcome Zn unavailability by taking the place of Zn in proteins whose function requires the presence of this metal cofactor (55). Previous observations have revealed that, also in *P. aeruginosa*, Co can repress the expression of Zur-dependent genes and stimulate the growth of a *znuAzrmA* mutant strain (19). Moreover, here we show that inactivation of the PCH biosynthetic pathway increases the demand for Zn, as assessed by the analysis of *zrmA* expression in the *znuAzrmB* and *znuAzrmBpchE* mutant strains (Fig 5, Panel A). A controlled increase in Co would therefore represent a strategy implemented by *P. aeruginosa* to cope with Zn deficiency conditions. However, it is necessary to highlight a significant difference in the responses of *Salmonella* Typhimurium and *P. aeruginosa* to Co. While high concentrations of Co have no obvious toxic effects on *Salmonella* (55), 10 µM Co inhibit the growth of *P. aeruginosa* (19). Results described in Figure 9 indicate that disruption of PCH synthesis ability increases resistance to Co of the *PA4065znuAzrmB* mutant strain, suggesting that Co toxicity could be at least partially correlated with intracellular accumulation of PCH. Therefore, PCH-mediated Co uptake pathway should be regarded as a double-edged strategy to face with Zn deficiency, as limited Co intake compensates for Zn deficiency, while excess Co is toxic. The need to carefully control the intracellular accumulation of Co can explain why an increase in the expression of the *zrmA* promoter is observed both in the *znuAzrmBpchE* strain (a strain that produces a very limited amount of PCH, as shown in Fig 5A) and in the *znuAzrmB* strain incubated with an excess of PCH (Fig 7). In fact, this latter condition could lead to an unnecessary intracellular Co accumulation. It is worth noting that many Zur-regulated genes with uncharacterized functions encode proteins annotated as proteins involved in cobalamin biosynthesis or import. Some of these proteins are putative metallochaperones of the COG0523 family, that are emerging as critical players in cell adaptation to conditions of limited Zn availability (56–58). It is tempting to hypothesize that these proteins play a role in the redistribution and allocation of Zn and Co in different proteins in response to a reduced intracellular availability of Zn.

The study also raises questions about the regulation of PCH synthesis, the selectivity of receptors for PCH-metal complexes, and the reasons behind changes in the extracellular accumulation of PCH. It is evident that, apart from *PA2911-2914*, representative genes involved in the synthesis (*pchD*, *pchE*) and transport of PCH-Fe (*fptA*, *fptX*) are strongly repressed in the *znuAzrmB* mutant. In contrast, PCH synthesis is active in *PA2914znuAzrmB* and *PA4065znuAzrmB* mutants. The repression of PCH synthesis in the *znuAzrmB* double mutant can probably be explained based on the model of PCH synthesis regulation recently proposed by Schalk and collaborators (51). According to this model, the transcription of all PCH genes is stimulated by the transcriptional activator PchR in complex with PCH–Fe^3+^ but is repressed by the binding of PchR to the PCH-Co complex. Under conditions of severe Zn deficiency that characterize the *znuAzrmB* mutant, increased expression of the *PA2911-2914* operon promotes the selective uptake of the PCH-Co complex, likely resulting in inhibition of PCH synthesis. Although FtpA can import PCH complexed with both Fe and Co, the uptake rate of Co is 26-fold lower than that for Fe (49). If we hypothesize that PA2911-2914 provides a more effective entry route for the PCH-Co complex, this would probably explain the fact that the inhibitory mechanism is not active in the *PA2914znuAzrmB* mutant. It will be interesting in the future to conduct studies to characterize the binding specificity of the FptA and PA2914 receptors in greater detail. However, the reason why PCH synthesis is reactivated in the *PA4065znuAzrmB* mutant despite an apparent intracellular accumulation of the PCH-Co complex remains to be investigated. For instance, it can be hypothesized that in the presence of high concentrations of this complex, the PchR regulator regains a transcriptionally active conformation.

Additionally, the identification of a MacB export system dedicated to PCH export requires some comment. While both the enzyme systems involved in the cytoplasmic synthesis of PCH and the transporters that mediate the entry of the PCH-Fe complex across the outer and inner membrane are well known, no protein complex involved in PCH secretion has ever been characterized (59). Given the predominantly hydrophobic nature of PCH, it has often been assumed that, once synthesized in the cytoplasm, it could permeate the membrane without a dedicated export system (60). Therefore, the MacB transporter, composed of the PA4064 and PA4065 subunits, is the first active PCH export system to be identified. This export system is under Zur regulation and is active only in conditions of Zn deficiency. Our results suggest that *P. aeruginosa* can export PCH even in the absence of this transporter. The mechanism for the export of newly synthesized PCH remains an open question, and membrane passage without a dedicated export system remains a valid hypothesis. Recent studies indicate that only a fraction of the PCH-Fe complex entering the periplasm via FptA is routed into the cytoplasm through FptX, while another part dissociates in the periplasm to allow free-form Fe entry through the channel formed by PchHI proteins (61). This is probably another mechanism to prevent intracellular accumulations of PCH that could be toxic to the cell. In this complex scenario, a diagram summarizing the role of the different characterized Zur-regulated metal transporters in *P. aeruginosa* response to Zn-deficiency is reported in Figure 10.

**Fig. 10.**
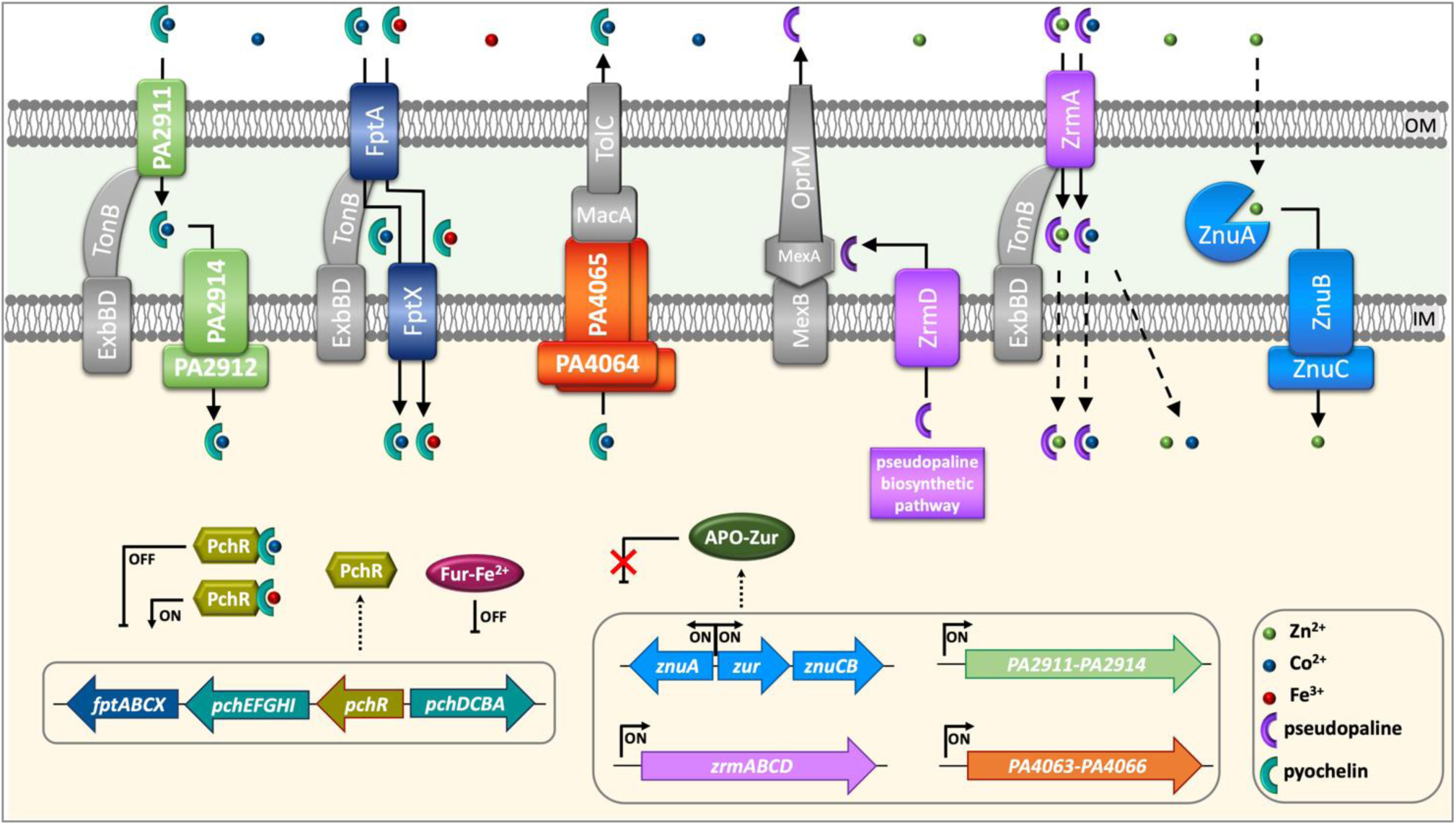
Schematic view of *P. aeruginosa* responses to Zn deficiency. The diagram illustrates the metal transport systems for which a role in the response to Zn deficiency is demonstrated. As in all Gram-negative bacteria, the Zur-regulated ZnuABC transporter has a central role in ensuring the transport of Zn from the periplasm to the cytoplasm (the dashed line through the Outer Membrane (OM) indicates that the entry routes of Zn into the periplasm are not defined). In addition to *znuABCD*, the operons *zrmABCD* (responsible for the synthesis and transport of pseudopaline), *PA2911-PA2914* and *PA4063-PA4066* are also negatively regulated by Zur under Zn-replete conditions and transcribed when the intracellular Zn concentration is low, a condition in which Zur is no longer properly metalated. The zincophore pseudopaline, synthesized in the cytoplasm by ZrmB and ZrmC, is transported into the periplasm by ZrmD and then exported out of the cell by the MexAB-OmpR efflux pump (62). Pseudopaline complexed to Zn or Co re-enters the periplasm via the TonB-dependent Outer Membrane Receptor ZrmA. It is currently unknown whether the pseudopaline-bound metal is released in the periplasm or transported into the cytoplasm in complex with the pseudopaline (dashed arrows through the Inner Membrane - IM). Co, which can partially compensate for conditions of severe Zn deficiency, can enter either in complex with pseudopaline or bound to PCH. The PCH-Co complex can enter the cell either from FptA, the receptor involved in the uptake of the PCH-Fe complex or through the PA2911-2914 import system. The binding of PCH-Co to PchR, the main regulator of PCH synthesis, represses the synthesis of PCH and of the FptA-FptX import system (51). In contrast, the expression of *PA2911-2914* is not repressed by PchR, but induced under conditions of poor Zn availability, thereby allowing the entry of PCH-Co even if *fptA-fptX* is repressed. To avoid potentially toxic accumulations of PCH and Co within the cell, excess PCH-Co is exported through a pump involving the MacB transporter formed by PA4064/PA4065.

Finally, the most intriguing aspect emerging from this study is a reevaluation of the paradigm regarding PCH as merely a Fe carrier. While *P. aeruginosa* can indeed produce pyoverdine, a siderophore with a significantly higher Fe affinity than PCH, it is now believed that under conditions of moderate Fe deficiency, PCH production is favored due to its lower energy requirements for synthesis compared to pyoverdine (63). This study unveils additional roles for PCH that extend beyond its conventional role in extracellular Fe uptake, as hinted at in a previous study (64). The findings reported here emphasize the involvement of PCH in cellular responses to Zn deficiency, shedding light on its broader role in connecting the homeostasis of Fe and Zn, two central metals in nutritional immunity. In this regard, several studies have highlighted that during infections many of the most highly expressed genes often belong to the Zur operon or are involved in the synthesis of PCH, suggesting that Fe and Zn are limited during infections (47,65). The identification of a role for PCH also in the adaptation of *P. aeruginosa* to Zn deficiency introduces new elements of complexity that suggest caution in the interpretation of transcriptomic data concerning the genes involved in the synthesis and transport of PCH.

In conclusion, our findings reveal that *P. aeruginosa* possesses intricate and finely tuned mechanisms for Zn import, explained by the critical role of Zn in infections. A comprehensive understanding of Zn acquisition systems can pave the way for new strategies to interfere with metal homeostasis, potentially enhancing infection control.

## Material and methods

### Reagents

Antibiotics, ethylenediaminetetraacetic acid disodium salt (EDTA), and 2-nitrophenyl β-D-galactopyranoside (ONPG) were acquired from Sigma-Aldrich (St. Louis, MO). Restriction endonucleases and T4 DNA ligase were purchased from New England Biolabs (Ipswich, MA). All other chemicals were purchased from VWR International (Milan, Italy). The oligonucleotides were synthesized by Eurofins Genomics (Milan, Italy) or Sigma-Aldrich (Darmstadt, Germany). Purified PCH was extracted from *P. aeruginosa* Δ*pvdA* (Table S1), quantified as previously described (66), and stored as a 40 mM stock solution in DMSO (Sigma-Aldrich, St. Louis, MO).

### Bacterial strains and growth media

The strains and plasmids used in this study are listed in Table S1. *E. coli* and *P. aeruginosa* strains were streaked respectively on Luria-Bertani Agar (LBA) (Bactotryptone 10 g L ^-1^, yeast extract 5 g L^-1^, 10 g NaCl 10 g L^-1^, BactoAgar 15 g L^-1^) or Pseudomonas Isolation Agar (PIA; Becton Dickinson, USA) plates. All the strains were routinely grown at 37 °C in LB-broth (LB). Zn-limiting growth conditions were achieved using the Vogel-Bonner Minimal Medium E (VBMM: MgSO_4_-7H_2_O 0.192 g L ^-1^, citric acid 2 g L ^-1^, K_2_HPO_4_ 10 g L ^-1^, NaNH_4_HPO_4_-4H_2_O 3.5 g L ^-1^, glucose 2 g L ^-1^), supplemented or not with 5 µM EDTA. VBMM was prepared in disposable plastic containers to minimize Zn contaminations and sterilized by filtration (19). Antibiotics were used at the following concentrations: for *E. coli* kanamycin 50 mg L^-1^, tetracycline 10 mg L^-1^, gentamicin 10 mg L^-1^ and carbenicillin 100 mg L^-1^; for *P. aeruginosa* gentamicin 100 mg L^-1^, tetracycline 100 mg L^-1^and carbenicillin 500 mg L^-1^.

### *P. aeruginosa* mutants construction

Mutant strains were obtained by the gene replacement method described by Hoang (67), following the same protocol previously used to generate *znuA*, *zrmA,* and *zrmB* mutants (19,21). The primers used for the generation of knock-out mutants are listed in Table S2. Cloning of the 5’ and the 3’ terminal fragments of the target genes and the gentamicin resistance cassette was performed in plasmid pEX18Tc, and the mobilization of the resulting vector from *E. coli* DH5α to *P. aeruginosa* was achieved by tri-parental mating, as already described (21). When needed, the gentamicin resistance cassette was removed from the strain using plasmid pFLP2, encoding the gene for the Flp recombinase, as previously described (21). Gene deletions or the loss of the gentamicin resistance cassette were confirmed by PCR using the primers listed in Table S2.

### Analyses of bacterial growth

Overnight cultures of *P. aeruginosa* grown in LB were diluted 1:1000 in VBMM with EDTA 5 μM (E-VBMM). A volume of 0.2 mL of each sample was inoculated in a 96-microwell (Greiner Bio-One, Austria) and incubated at 37°C in a Sunrise™ microplate reader (Tecan, Männedorf, Switzerland). Optical density at 595 nm (OD_595_) was registered every hour for 24 hours. Each sample was tested in triplicate and the experiment was performed at least two times with comparable results.

### RNA extraction and Real Time-PCR analysis

RNA was extracted from *P. aeruginosa* PA14 using the RNAeasy kit (Qiagen, Hilden, Germany) according to the manufacturer’s protocol, with the addition of DNase (Qiagen) and lysozyme (Sigma Aldrich). The concentration and purity of the RNA were determined with a NanoDrop™ Lite Spectrophotometer (Thermo Fisher Scientific). Three replicates for each experimental condition were prepared. cDNA was synthesized from 1 µg of each RNA sample using PrimeScript RT Reagent Kit with gDNA Eraser (Takara), and the primers used for qRT-PCR were designed using Primer3web, version 4.1.0 and are listed in Table S2. qRT-PCR reactions were performed in 10 µL mixtures containing 50 ng cDNA, 0.3 µM of each primer, and 50% SYBR green (PowerUp SYBR Green Master Mix, Thermo Fisher, Waltham MA), using a Thermo Fisher (QuantStudio3) thermocycler with the following parameters: (i) initial denaturation, 4 min at 95°C; (ii) 40 cycles of denaturation, primer annealing, and extension (20 s at 95°C, 30 s at 60°C, 30 s at 72 °C ); (iii) production of melting curve, from 50°C to 90°C (rate: 0.58°C every 5 s). The relative amount of mRNA of each gene was determined using the 2^-ΔΔCt^ formula, where the threshold cycle (Ct) of the target gene in the treated sample is normalized to the reference gene (*oprI* or *rpoD*) and the respective value obtained in the untreated bacteria. Each experiment was repeated at least twice, obtaining comparable results.

### Fractionation of *P. aeruginosa* supernatants

Bacteria were grown in 20 ml of VBMM for 18 hours at 37°C under shaking. Cells were centrifuged at 5000 rpm for 15 min, and supernatants were collected and filtrated using a 0.22 µm syringe filter (Cytiva, Marlborough, MA, USA). The filtrates were acidified with 37% HCl to pH 2.0-3.0, diluted with 0.4 volumes of ethyl acetate, and centrifuged at 5000 rpm for 15 min. Ethyl acetate addition and centrifugation were repeated three times. The organic phases were collected and dried in SC110 Speedvac® Concentrator (Savant Instruments, Holbrook, NY) for up to three hours. The resulting pellets were resuspended 1:20 (vol:vol) methanol:ddH_2_O for the feeding assays and 1:1 (vol:vol) methanol:ddH_2_O for spectrophotometric analysis.

### *zrmA* promoter activity assay

The reporter plasmid *pzrmA*pTZ110 (19), carrying the promoter region of *zrmA* upstream of the *lacZ* gene, was mobilized from *E. coli* DH5α into PA14 strains by triparental mating, using *E. coli* HB101 pRK2013 as the helper strain (21). PA14 exconjugants carrying *pzrmA*pTZ110 were pre-inoculated overnight in LB medium, diluted 1:1000 in E-VBMM, and incubated at 37°C for 18 hours. For each experimental condition, three independent colonies were tested, following the beta-galactosidase activity assay previously described (23). In triplicate, absorbances were measured in Sunrise™ reader (Tecan, Männedorf, Switzerland).

### Modeling of PA4064/PA4065 and PA2911-PA2914

The structures of the individual proteins encoded by the *P. aeruginosa PA2911-PA2914* gene cluster (UniProt ID: Q9HZT6, Q9HZT5, Q9HZT4, and Q9HZT3) were modeled through the SWISS-MODEL web server (68). The model of the uncharacterized complex PA4064-4065 (UniProt ID: Q9HWW3 and Q9HWW4) was obtained by the Alphafold 2 method (69), applying the AlphaFold-multimer prediction model, using the full database for the construction of multiple sequence alignments (MSAs) and excluding from the pipeline the 3D structures published before December 31^st^, 2000. In this technique, a folded protein is considered a “spatial graph” where residues are the nodes, and edges connect the residues in close proximity. An attention-based neural network system, trained end-to-end, attempts to interpret the structure of this graph while reasoning over the implicit graph has been created in the latest version of AlphaFold used at CASP14 (Critical Assessment of Techniques for Protein Structure Prediction, May-August 2020).

### Metal analysis

*P. aeruginosa* strains were grown in 10 mL of E-VBMM containing trace amounts of transition metals, namely ZnSO_4_ 0.2 μM, FeSO_4_ 0.1 μM, NiSO_4_ 0.1 μM, Co(NO_3_)_2_ 0.1 μM, CuSO_4_ 0.1 μM, and MnCl_2_ 0.1 μM. Cultures were grown for 18 h at 37°C in 50 mL polypropylene tubes, collected, and pelleted by centrifugation at 6000 rpm for 10 min. The samples were carefully washed with 10 mL of phosphate buffer saline (PBS) containing EDTA 1 mM to remove metals weakly bound to the bacterial surface. Bacterial pellets were dried in SC110 Speedvac® Concentrator and stored at -20°C for further analyses. Metal content in bacterial pellets was determined by an Inductively Coupled Plasma Mass Spectrometer (ICP-MS, model 820-MS; Bruker, Bremen, Germany) equipped with a collision–reaction interface (CRI). Fe and Mn were analyzed using ICP-MS in CRI mode with H_2_ and He (99.9995% purity; SOL Spa, Monza, Italy) as cell gases; the other elements by ICP-MS in standard mode. The ICP-MS optimized instrumental parameters are summarized in a previous study (70). Multi-element standard solutions (VWR International, Milan, Italy) were used for instrumental calibration curves (seven-point). Yttrium at 0.005 mg L^−1^ (Panreac Química, Barcelona, Spain) and Scandium, and Indium at 10 mg L^−1^ (Merck, Darmstadt, Germany) were used as internal standards. Digestion of the bacterial pellet samples (∼10 mg; three replicates) was performed in a water bath (WB12; Argo Lab, Modena, Italy) using a mixture of 0.2 mL HNO_3_ and 0.1 mL H_2_O_2_, according to (71). The obtained digests were diluted to 5 mL with deionized water (18.3 MΩcm resistivity) obtained from an Arioso (Human Corporation, Seoul, Korea) Power I RO-UP Scholar UV deionizer system, and filtered (0.45μm pore size, GVS Filter Technology, Indianapolis, IN, United States). Method blanks were periodically analyzed alongside the samples to check for losses or cross-contamination. The method detection and quantification limits (MDL and MQL, respectively) were 0.009 and 0.03 mg kg^−1^ for Co, 4 and 10 mg kg^−1^ for Fe, 0.05 and 0.2 mg kg^−1^ for Mn, 0.2 and 0.5 mg kg^−1^ for Ni, and 0.8 and 3 mg kg^−1^ for Zn, respectively.

### PCH quantification in cell lysates and culture supernatants

Bacteria grown overnight at 37°C in 10 ml of VBMM were collected, and the OD_600_ was recorded. The cultures were centrifuged at 6000 rpm for 10 min. The supernatants were filtered and stored at - 20°C. The pellet of each sample was resuspended in a sonication buffer (MgSO_4_-7H_2_O 0.192 g L^-1^, anhydrous K_2_HPO_4_ 10 g L^-1^, NaNH_4_HPO_4_-4H_2_O 3.5 g L^-1^) at a final concentration of 20 OD_600_ ml^-1^. Cells were sonicated using a Sonifier® Cell Disruptor (Branson Ultrasonics Corporation, Danbury, CT) in an ice bath at 25% amplitude for 15 min.

PCH detection in culture supernatants and cell lysates was performed using the *P. aeruginosa ΔpvdAΔpchDΔfpvA PpchE*::*lux* biosensor as previously described (35). Culture supernatants and cell lysates were diluted at 1:250 and 1:50 in the Fe-poor casamino acid medium DCAA (25). Five microliters of appropriate dilutions of culture supernatants or cell lysates were added to 45 μL of DCAA inoculated with the biosensor strain (final OD_600_ = 0.25). Microtiter plates were incubated at 25 °C, and OD_600_ and LCPS were measured after 3.5 h using a Tecan Spark 10 M multilabel plate reader. A calibration curve was generated with purified PCH at known concentrations (5−320 nM) and used to calculate the concentration of PCH in each sample.

## Acknowledgments

The authors thank Prof Luigi Scipione (University of Rome Sapienza) for the MS analysis of the supernatant fractions extracted with ethyl acetate. This study was partially supported by grants to AB from Cystic Fibrosis Foundation (BATTIS19IO) and Italian MUR (PRIN 2022, contract 2022E57Z3K, CUP E53D23009850006, Funded by the European Union – NextGenerationEU”). AB and PV were also supported by Project ECS 0000024 Rome Technopole, – CUP B83C22002820006, NRP Mission 4 Component 2 Investment 1.5, Funded by the European Union – NextGenerationEU”.

