## Supplementary Figures and Tables for "Investigation of Zur-regulated metal transport systems reveals an unexpected role of pyochelin in zinc homeostasis"

**S1 Fig.**

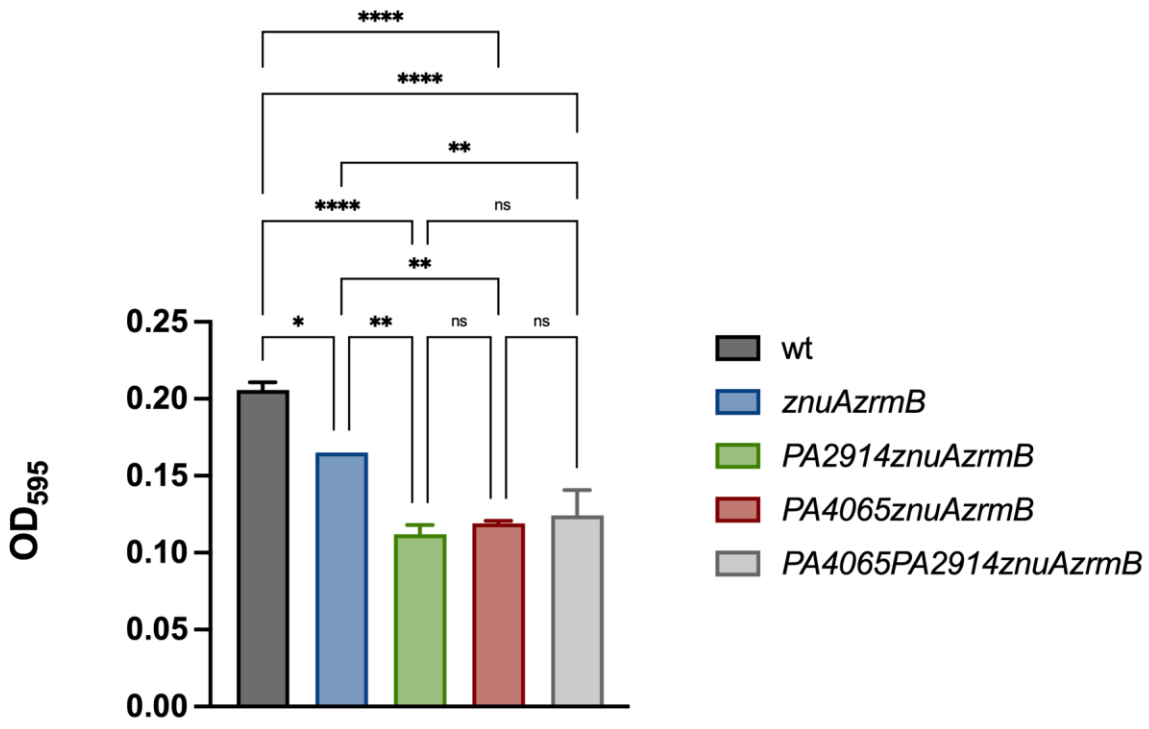


**The simultaneous deletion of *PA4065* and *PA2914* in the *znuAzrmB* background does not cause any growth significant difference compared to *PA2914znuAzrmB PA4065znuAzrmB* and triple mutants.** Bars represent mean values with SD of three independent cultures. Statistical significances were calculated by two-way ANOVA and Bonferroni's multiple comparison test. Asterisks indicate statistically significant differences (*p<0.05; **p<0.005; ****p < 0.0001).

**S2Fig.**

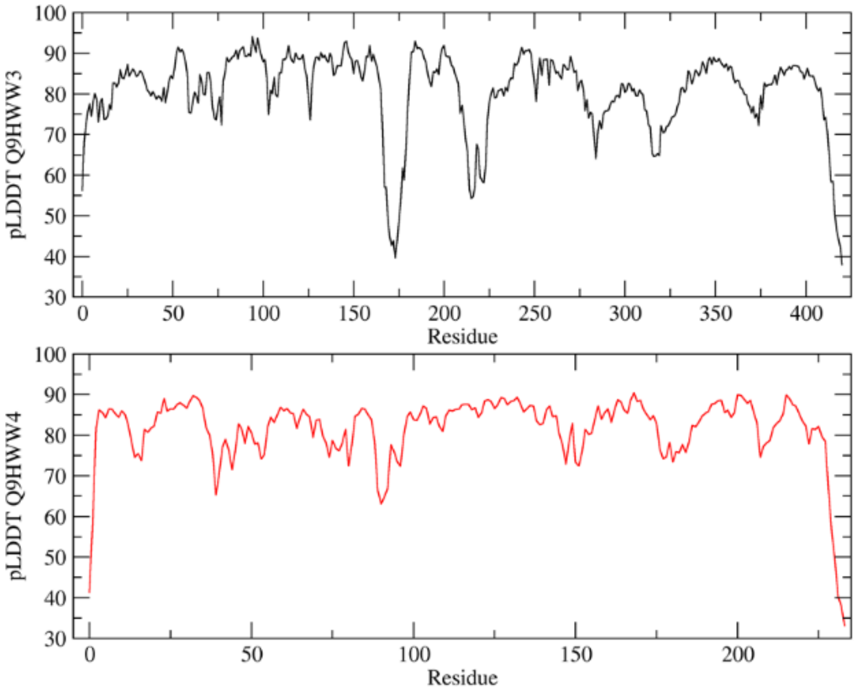


**pLDDT values.** Values are obtained as a function of the sequence observed for the structures predicted by Alphafold 2

### S3 Fig.

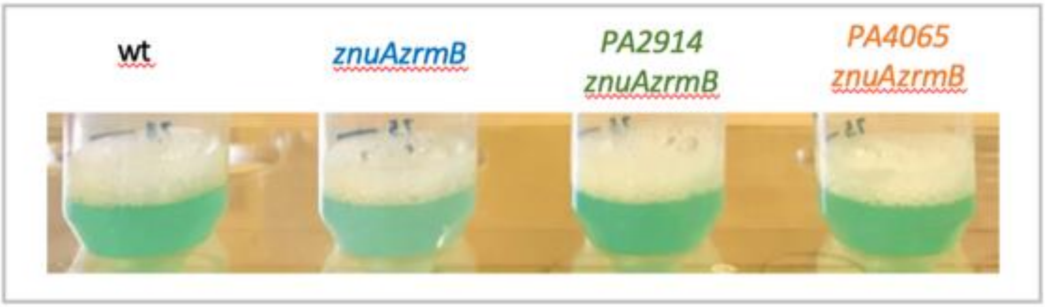


**Pigmentation of stationary phase cultures of wild-type PA14 and mutant strains *znuAzrmB*, *PA4065znuAzrmB*, and *PA2914znuAzrmB*, grown in VBMM**. The *znuAzrmB* strain shows a different pigmentation compared to the other strains

### S4 Fig.

#### A) PAO1 wild type

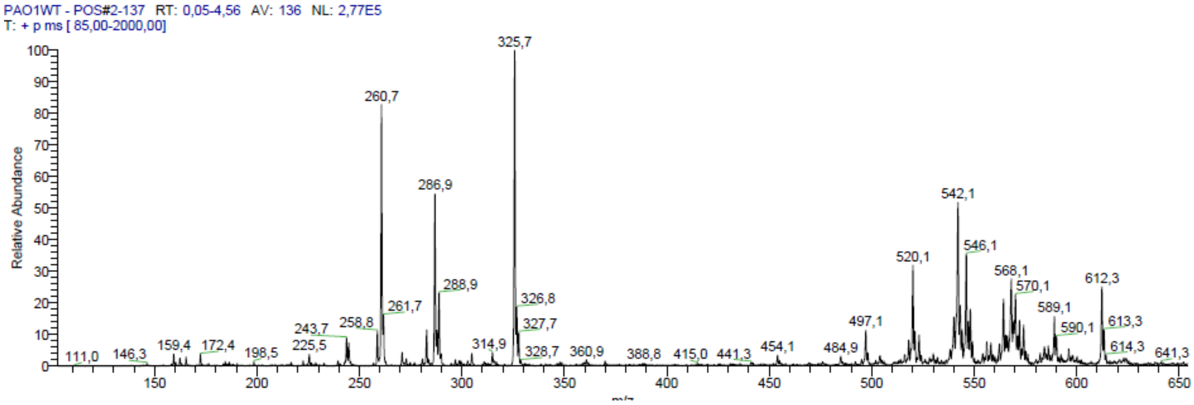


#### B) PAO1 *pchD*

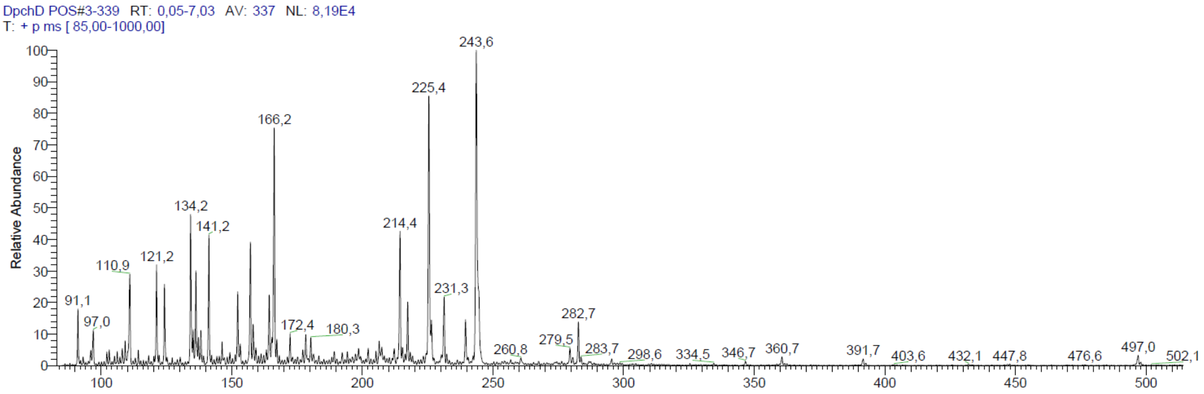


#### C) PA14 wild type

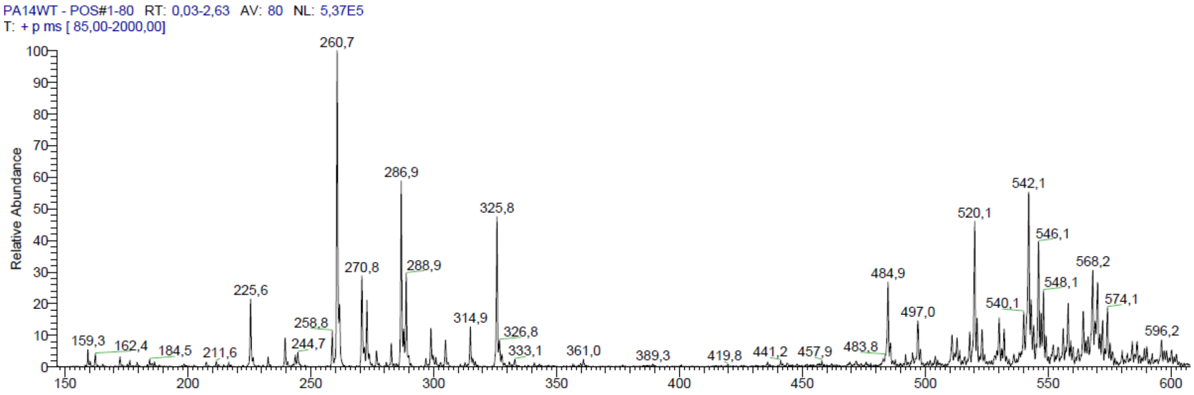

**D) PA14 *znuAzrmB***

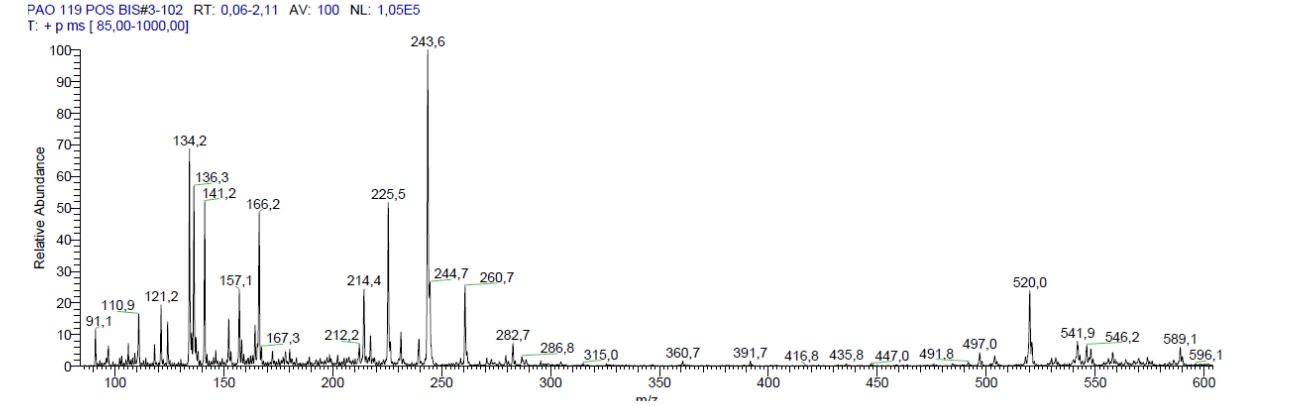


**E) PA14 *PA2914znuAzrmB***

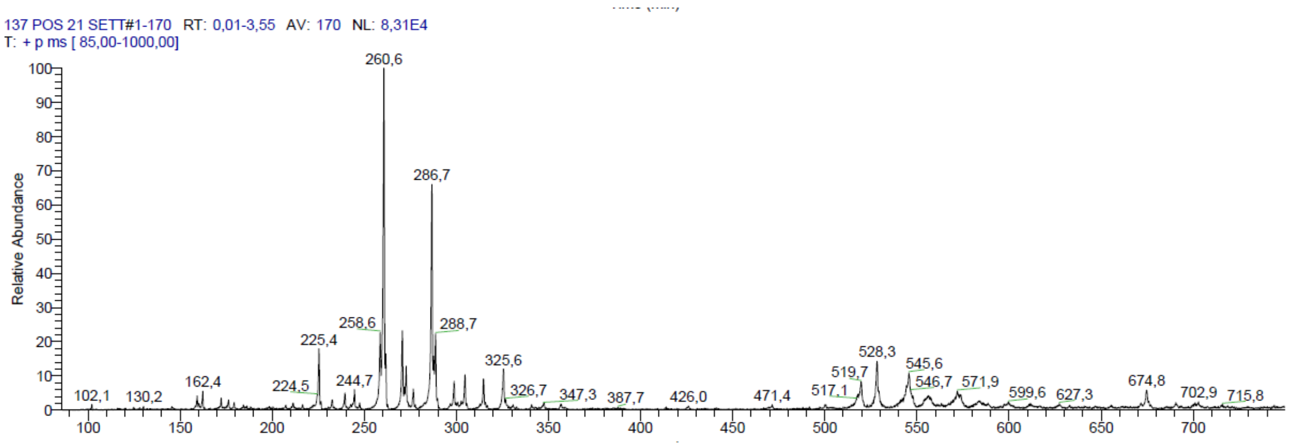


**F) PA14 *PA4065znuAzrmB***

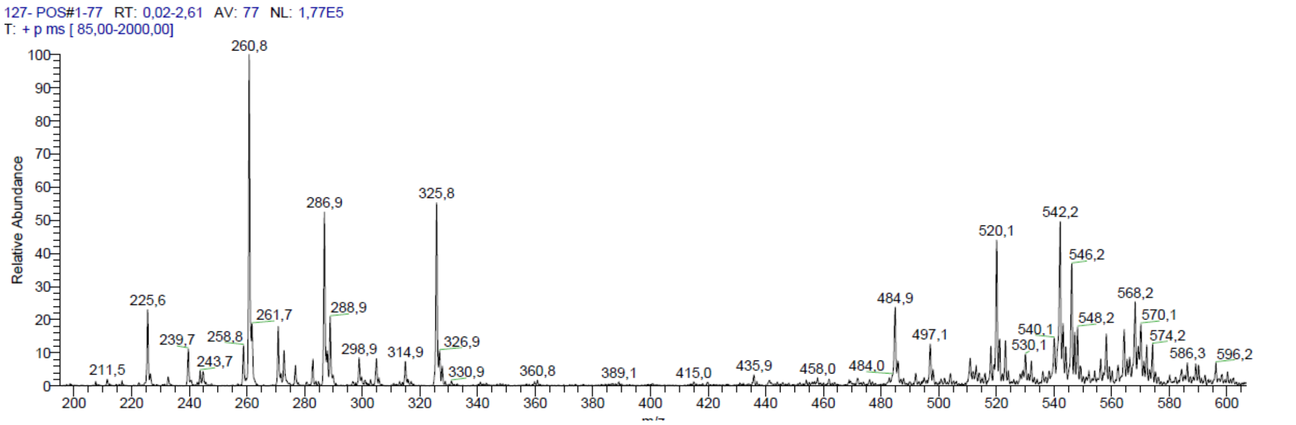


**Positive mode ESI spectra of the supernatants (fraction extracted with ethyl acetate) from *Pseudomonas aeruginosa* PAO1 and PA14.**

The mass spectra were recorded on a ThermoFinnigan LCQ Classic LC/MS/MS ion trap equipped with an ESI source and an injection syringe pump. The fractions of the supernatants extracted with ethyl acetate were diluted 1/10 with MeOH/$H_2O$ (9:1) and were infused in the electrospray system at a constant flow rate of 10 µL/min$^{-1}$ and the mass spectra were recorded in positive ion mode (spray voltage 6.5kV, capillary voltage 35V, capillary temperature 200°C). ESI-MS data are given as m/z, with mass expressed in atomic mass units (amu).

Panels A and B compare the fractions extracted from the wt PAO1 strain and the *pchD* strain. The spectrum of the sample from the *pchD* strain shows an extremely simplified profile compared to that of the wild strain. In addition to missing the peak corresponding to PCH (m/z 325), all other major peaks are absent in the mutant, including the one with m/z 260 (PQS), the peaks with m/z 286 and 288, likely corresponding to other 4-hydroxy-2-alkylquinolines (HAQs)(1) and all those with m/z above 500, possibly corresponding to rhamnolipids (2). The peaks observed in the spectrum of the mutant represent molecular species that are very scarcely abundant in the wild-type strain. Similarly, the main ion species present in the PA14 wild type strain supernatant (panel C) are absent in the *znuAzrmB* mutant (panel D), whose composition is similar to that of the *pchD* mutant reported in panel B (absence of peaks for PCH and PQS). In contrast, the mass spectrum of the *PA4065znuAzrmB* mutant (panel F) is very similar to that of the wild-type strain. The peaks corresponding to the various HAQs and PCH are also restored in the *PA2914znuAzrmB* triple mutant, although in this case, the peak for PCH is less pronounced than those of HAQs

### S5 Fig.

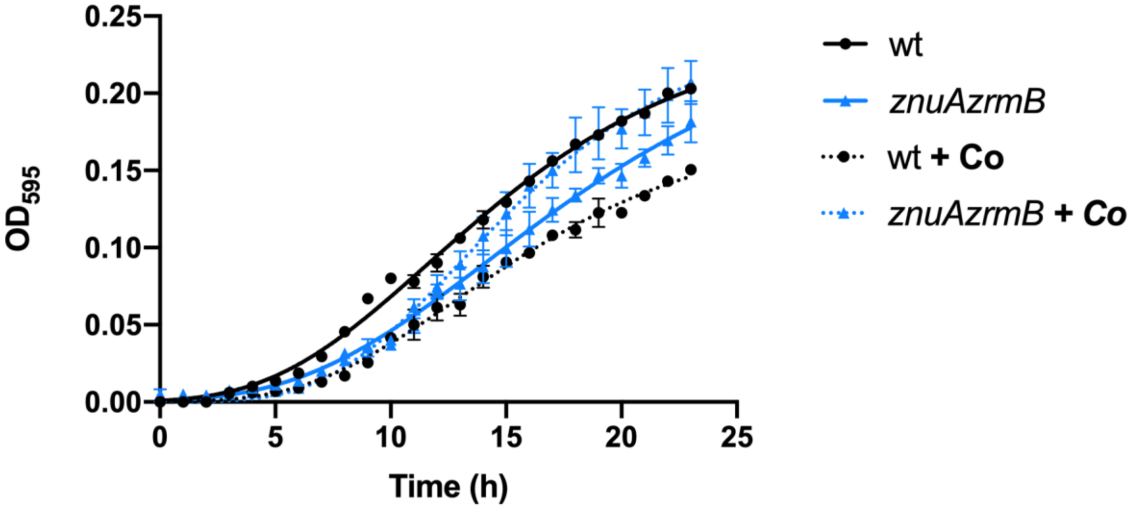


**Effect of Co supplementation on PA14 wild type and *znuAzrmB* strains.** Bacteria were grown in E-VBMM (continuous lines) and E-VBMM + $Co(NO_3)_2$ 10 µM (dotted lines). Each symbol indicates the mean ± SD of triplicates, and lines represent nonlinear fit according to the Logistic Growth equation.

**Table S1: Bacterial strains and plasmids used in this work.**

| *Strain* | *Relevant genotype* | *Reference or source* |
|---|---|---|
| ***E. coli*** | | |
| DH5α | φ80 Δ*lacZ* 15Δ (*lac-arg*F) U169 *deo*R*rec*A1 *end*A1 *hsd*R17 ($rk^-$, $mk^+$) *pho*A*sup*E44 λ$^-$*thi*-1 *gyr*A96 *rel*A1 | Lab collection |
| HB101 | $F^-$ *mcrB mrr hsdS20*($r_B^-$ $m_B^-$) *rec*A13 *leu*B6 *ara*-14 *pro*A2 *lacY1 gal*K2 xyl-5 mtl-1 *rps*L20($Sm^R$) glnV44 λ$^-$ | Lab collection |
| ***P. aeruginosa*** | | |
| PA14 | wild type | Lab collection |
| PA14_MDO101 | *znuA::Gm* | (3) |
| PA14_MDO111 | *zrmA::Gm* | (4) |
| PA14_MDO113 | *znuA::scar zrmA::Gm* | (4) |
| PA14_MDO117 | *zrmB::Gm* | This study |
| PA14_MDO118 | *PA4065::Gm* | This study |
| PA14_VS-MDO127 | *PA4065::scar znuA::scar zrmB::Gm* | This study |
| PA14_VS-MDO129 | *PA2914::Gm* | This study |
| PA14_VS-MDO137 | *znuA::scar PA2914::scar zrmB::Gm* | This study |
| PA14_VS-MDO152 | *PA4065::scar znuA::scar zrmB::scar PA2914::Gm* | This study |
| PA14_VS-MDO153 | *pchE::Gm* | This study |
| PA14_VS-MDO154 | *znuA::scar zrmB::scar pchE::Gm* | This study |
| PA14_VS-MDO155 | *znuA::scar zrmB::scar PA2914::scar pchE::Gm* | This study |
| PA14_VS-MDO156 | *znuA::scar zrmB::scar PA4065::scar pchE::Gm* | This study |
| PAO1 | wild type | Lab collection |
| PAO1 | *ΔpchD* | Lab collection |
| PAO1 | *ΔpvdAΔpchDΔfpvA PpchE::lux* | (5) |
| PAO1 | *ΔpvdA* | (6) |
| ***Plasmids*** | ***Description*** | ***Reference*** |
| pEX18Tc | Broad-host-range gene replacement vector with MCS from pUC18; *sacB*+;TcR,*oriT*+ | (7) |
| pRK2013 | Broad-host-range helper vector; *ColE1*- | (8) |
| pFLP2 | BhrFlp recombinase-producing plasmid | (7) |
| p*zrmA*PTZ110 | Reporter plasmid bearing a *zrmA* promoter::*lacZ* fusion | (4) |
| pPS856 | Source of gentamicin resistance cassette. AmpR, GmR | (7) |

**Table S2: List of primers used in this work.**

| **Primers for mutant strains construction:** | | |
|---|---|---|
| **Primer** | **Sequence (5'-3')** | **Description** |
| PA2914_1 | ATAGAATTCGTGTTCCTCTACGACAGC | Forward primer to amplify the 5' region of *PA2914* |
| PA2914_2 | ATAGGATCCGATCAGCCAGACGATGTG | Reverse primer to amplify the 5' region of *PA914* |
| PA2914_3 | ATAGGATCCCTGTGGGTCTTCATCTGC | Forward primer to amplify the 3' region of *PA2914* |
| PA2914_4 | ATAAAGCTTTGTTCATGCCCGACTACG | Reverse primer to amplify the 3' region of *PA2914* |
| PA2914_5 | *TCATCGTCATCGTCGACTAC* | Forward primer to check *PA2914* deletion |
| PA2914_6 | *TTCATGTGCCACGACTACAC* | Reverse primer to check *PA2914* deletion |
| PA4065_5 | CTCGAATTCGGCGTTTCTCCAGTTGCTCT | Forward primer to amplify the 5' region of *PA4065* |
| PA4065_6 | CGCGGATCCCTTGTCGTCGTGCTTGACCA | Reverse primer to amplify the 5' region of *PA4065* |
| PA4065_7 | CGCGGATCCCAGGCCAACTACGGCATCTA | Forward primer to amplify the 3' region of *PA4065* |
| PA4065_8 | CCCAAGCTTCTGGCGTTCTCCACCTTGAA | Reverse primer to amplify the 3' region of *PA4065* |
| PA4226_9 | CCCGAGCTCATCTAACGAAACAGTCCG | Forward primer to amplify the 5' region of *pchE* |
| PA4226_2 | ATAGGATCCAACATCGGGTGGCGTTGA | Reverse primer to amplify the 5' region of *pchE* |
| PA4226_3 | ATAGGATCCGACCGGGTGATCAGCTTC | Forward primer to amplify the 3' region of *pchE* |
| PA4226_4 | CTCAAGCTTCTCCAGTTGGGTTTCCTC | Reverse primer to amplify the 3' region of *pchE* |
| PA4226_7 | *ATCCGCCTGATGTACCTGCA* | Forward primer to check *pchE* deletion |
| PA4226_8 | *ATCGACCTTGCCATTGCC* | Reverse primer to check *pchE* deletion |
| **Primers for qPCR:** | | |
| **Target gene** | **Forward (5'-3')** | **Reverse (5'-3')** |
| *fptA* | ACCTACGAAACCGGGATCAA | GGTCTTCCTGCGGATTGTTC |
| *fptX* | CTGGGTGGTCAAGTTCCTCT | ATCGGCAGGATCCAGCTAC |
| *oprI* | ATTCTCTGCTCTGGCTCTGG | CGGTCTGCTGAGCTTTCTG |
| *PA2911* | CTACATCGACCCCTGGCATC | GCGGTCGATATGGTTCTGGT |
| *PA4063* | CACAAGGAGAAGGCAGGCC | CTGAATTTTCTGGGTGGCGG |
| *pchD* | CCTTCGTCGAGACCTGCTT | CTGATCTCATGCTGGCGATG |
| *pchE* | GATCAATACCATCGACGCGC | GATAGATCGAAGTCCAGCGC |
| *pchR* | CATCACCATCATTGCTCCGC | GGTCACCAGCTTCATATTCGG |
| *rpoD* | CATCGCCAAGAAGTACACCA | CCACGACGGTATTCGAACTT |

**Table S3: ICP-MS analyses of intracellular Fe content.**

| Strain | Fe56 (mg/kg) |
|---|---|
| Wild type | 88.0 ± 3.2 |
| *znuA* | 90.0 ± 2.6 |
| *pchE* | 77.0 ± 2.2 |
| *znuAzrmB* | 84.0 ± 1.7 |
| *znuAzrmBpchE* | 83.0 ± 3.3 |
| *PA2914znuAzrmB* | 84.0 ± 13.3 |
| *PA2914znuAzrmBpchE* | 78.0 ± 0.7 |
| *PA4065znuAzrmB* | 90.0 ± 9.2 |
| *PA4065znuAzrmBpchE* | 76.0 ± 1.9 |

**Supplementary Figures References**
